# Sources of variation in the spectral slope of the sleep EEG

**DOI:** 10.1101/2021.11.08.467763

**Authors:** N Kozhemiako, D Mylonas, JQ Pan, MJ Prerau, S Redline, SM Purcell

## Abstract

Building on previous work linking changes in the electroencephalogram (EEG) spectral slope to arousal level, Lendner et al. (2021) reported that wake, non rapid eye movement (NREM) sleep and rapid eye movement (REM) sleep exhibit progressively steeper 30-45 Hz slopes, interpreted in terms of increasing cortical inhibition. Here we sought to replicate Lendner et al.’s scalp EEG findings (based on 20 individuals) in a larger sample of 11,630 individuals from multiple cohorts in the National Sleep Research Resource (NSRR). In a final analytic sample of *N* = 10,255 distinct recordings, there was unambiguous statistical support for the hypothesis that, within individuals, the mean spectral slope grows steeper going from wake to NREM to REM sleep. We found that the choice of mastoid referencing scheme modulated the extent to which electromyogenic or electrocardiographic artifacts were likely to bias 30-45 Hz slope estimates, as well as other sources of technical, device-specific bias. Nonetheless, within individuals, slope estimates were relatively stable over time. Both cross-sectionally and longitudinal, slopes tended to become shallower with increasing age, particularly for REM sleep; males tended to show flatter slopes than females across all states. Although conceptually distinct, spectral slope did not predict sleep state substantially better than other summaries of the high frequency EEG power spectrum (>20 Hz, in this context) including beta band power, however. Finally, to more fully describe sources of variation in the spectral slope and its relationship to other sleep parameters, we quantified state-dependent differences in the variances (both within and between individuals) of spectral slope, power and interhemispheric coherence, as well as their covariances. In contrast to the common conception of the REM EEG as relatively wake-like (i.e. ‘paradoxical’ sleep), REM and wake were the most divergent states for multiple metrics, with NREM exhibiting intermediate profiles. Under a simplified modelling framework, changes in spectral slope could not, by themselves, fully account for the observed differences between states, if assuming a strict power law model. Although the spectral slope is an appealing, theoretically inspired parameterization of the sleep EEG, here we underscore some practical considerations that should be borne in mind when applying it in diverse datasets. Future work will be needed to fully characterize state-dependent changes in the aperiodic portions of the EEG power spectra, which appear to be consistent with, albeit not fully explained by, changes in the spectral slope.

## Introduction

Building on prior work^1–5^, Lendner et al. (ref ^6^) recently reported an electrophysiological marker of arousal level in humans: the 1/*f* spectral slope of the electroencephalogram (EEG) estimated in the range of 30 to 45 Hz. When calculated so as to avoid frequencies with strong oscillatory components (such as spindle activity during N2 sleep), the linear slope of the log-log power spectrum (the so-called ‘spectral noise’ exponent) can be interpreted as an index of the aperiodic, scale-free component that reflects aggregated neural dynamics^7^. Under this model, more synchronized activity results in steeper (more negative) 1/*f* slopes ^7–10^. Furthermore, it has been suggested that the spectral slope might reflect the ratio of neural excitation and inhibition (E/I balance)^2, 11, 12^. Lendner et al. found significantly steeper slopes (consistent with greater cortical inhibition) in REM compared to NREM sleep, and in NREM sleep compared to wake, concluding that the spectral slope represents a biomarker of human arousal, that might have applications in intraoperative neuromonitoring or automatic sleep stage classification, for example.

Here, we initially sought to replicate Lendner et al.’s core result concerning scalp EEG spectral slope and sleep state, leveraging a collection of over 15,000 whole-night polysomnograms (PSGs) from the National Sleep Research Resource (NSRR). Whereas Lendner et al.’s analyses were restricted to a small sample (*N* = 20 for the primary scalp sleep EEG dataset) albeit one complemented with additional imaging and intracranial EEG recordings, our analyses of multiple, diverse cohorts were intended to provide high statistical power, robustness and generalizability across populations, albeit based on only two central scalp EEG channels available across all studies. (Lendner et al. reported broad effects across the scalp including central sites, and source localization is not a focus of this report.) To address its potential role in automated sleep staging, we also asked whether the spectral slope predicted sleep state (epoch by epoch) with greater accuracy than traditional (e.g. band power) metrics.

Lendner et al. employed various checks for potential methodological confounds, including referencing scheme and analytic approach. Although they did not look at contralateral mastoid referencing (i.e. in the present context, C4-M1 and C3-M2), slope estimates were broadly consistent across the different referencing schemes considered (linked mastoid, average, Laplacian and bipolar). We initially adopted contralateral mastoid referencing, which is recommended for clinical studies by the American Academy of Sleep Medicine (AASM); further, the SHHS dataset (total *N* = 8,444 nights on 5,793 individuals) was recorded with contralateral references hardwired and so could not be re-referenced offline. Lendner et al. briefly considered potential confounding due to muscle activity: controlling for the spectral slope derived from the chin EMG, they reported that this did not alter results. Nonetheless, that the electromyogram (EMG) is 1) of orders of magnitude higher amplitude than the EEG, 2) exhibits a broad frequency spectrum that overlaps the EEG spectrum, particularly at higher (>30 Hz) frequencies ^13^ and 3) varies markedly between wake, NREM and REM ^14^, collectively make this a serious concern that should always be addressed. In particular, as mastoid electrodes are sensitive to neck muscle EMG and cardiac activity (via blood flow in the carotid arteries), here we considered possible indicators of confounding due to non-neural sources, with attention to the choice of mastoid referencing scheme.

Beyond our primary replication attempt - to robustly establish mean slope differences between states - in order to more fully characterize spectral slope distributions we evaluated its variability as a function of sleep state, as well as state-specific covariation between spectral slope, power and coherence. Using model-based simulation, we considered whether changes in the spectral slope alone (assuming a strict power law model) could account for these other characteristics.

Finally, as well as within-individual, between-state phenomena, we investigated between-individual, within-state changes in the spectral slope. Previous reports have suggested that the spectral slope varies between individuals in systematic and physiologically relevant ways, for example, flattening with age ^15–20^. As NSRR cohorts included males and females from ~5 to ~97 years of age (and a sample of individuals with a repeated sleep study, performed years later), we also tested whether the spectral slope showed age-related flattening and sex differences, and whether these effects varied by sleep state.

## Results

We analyzed 15,709 whole-night polysomnograms (PSGs) on 11,630 individuals (**Table 1**), 4,079 of whom had a second PSG, from the National Sleep Research Resource (NSRR). This is the same sample (comprising ten cohorts from six distinct studies) as previously described in a study of sleep spindle activity ^21^. Three cohorts were pediatric, six were of middle- to late-adulthood, and one was a family-based study with a wide age range. Other than CHAT, all cohorts were observational and not undergoing sleep or other interventions.

**Table 1:** Cohort characteristics. All data are available via the National Sleep Research Resource (http://sleepdata.org). CHAT(FU), SHHS2 and MrOS2 cohorts contained repeated PSGs performed on subsets of CHAT(BL), SHHS1 and MrOS1.

| Study | Label | Sample N | % female | Mean age | Min. age | Max. age | DOI |
| --- | --- | --- | --- | --- | --- | --- | --- |
| Childhood Adenotonsillectomy Trial (baseline) | CHAT(BL) | 453 | 52% | 6.6 | 5 | 10 | https://doi.org/10.25822/d68d-8g03 |
| Childhood Adenotonsillectomy Trial (non-randomized) | CHAT(NR) | 779 | 53% | 7.1 | 5 | 10 | https://doi.org/10.25822/d68d-8g03 |
| Childhood Adenotonsillectomy Trial (follow-up) | CHAT(FU) | 407 | 51% | 7.1 | 5 | 10 | https://doi.org/10.25822/d68d-8g03 |
| Cleveland Children's Sleep and Health Study | CCSHS | 515 | 50% | 17.7 | 16 | 20 | https://doi.org/10.25822/cg2n-4y91 |
| Cleveland Family Study | CFS | 730 | 55% | 41.4 | 7 | 89 | https://doi.org/10.25822/jmyx-mz90 |
| Sleep Heart Health Study (wave 1) | SHHS1 | 5793 | 52% | 63.1 | 39 | 90 | https://doi.org/10.25822/ghy8-ks59 |
| Sleep Heart Health Study (wave 2) | SHHS2 | 2647 | 54% | 67.6 | 44 | 90 | https://doi.org/10.25822/ghy8-ks59 |
| Osteoporotic Fractures in Men Study (wave 1) | MrOS1 | 2907 | 0% | 76.4 | 67 | 96 | https://doi.org/10.25822/kc27-0425 |
| Osteoporotic Fractures in Men Study (wave 2) | MrOS2 | 1025 | 0% | 81.1 | 73 | 97 | https://doi.org/10.25822/kc27-0425 |
| Study of Osteoporotic Fractures | SOF | 453 | 100% | 82.9 | 75 | 95 | https://doi.org/10.25822/e1cf-rx65 |

After stringent exclusions and quality controls (QC, see **Methods** & **Table S1**), the final analytic sample comprised 10,255 nights on 7,312 individuals. Most primary analyses were based on the first recording from each individual, pooling CHAT baseline and non-randomized samples, yielding six independent cohorts. Cohorts containing repeated recordings (CHAT follow-up, SHHS2 and MrOS2) were used in the longitudinal analyses, paired with their respective baseline cohort. All analyses used our open source Luna package (http://zzz.bwh.harvard.edu/luna/), with the main script used presented below, **Supplementary Information, Section 3.**

### Initial analyses of spectral power and slope stratified by sleep state

Primary analyses were based on a three-level classification: wake, N2 sleep and REM sleep. As more individuals had a sufficient duration of N2 sleep following extensive QC (at least 10 epochs), compared to N1 or N3 (**Table S1**), we focussed specifically on N2 sleep for all primary NREM analyses. (As results for N1 and N3 were broadly equivalent to those for N2, Figures and Tables only show N2 results but refer to it as ‘NREM sleep’ generically.) We first estimated each study’s mean log-scaled EEG power spectra (C4-M1) during wake, NREM and REM (**Figure 1a**). Perhaps most notably, both SHHS studies (blue lines) showed divergent, supralinear mean spectra across all states, especially >30 Hz. On further investigation, we identified a set of technical issues specific to the SHHS that impacted the high frequency EEG (**Figure S1** & **Supplementary Information Section 1**). Beyond this and as expected, wake power spectra exhibited characteristic ~8 Hz alpha peaks across all cohorts, whereas N2 spectra exhibited characteristic sigma-band peaks. The childhood CHAT samples (green lines) had more pronounced spindle peaks at slower frequencies (e.g. 11 Hz) and, across all states, higher power at lower frequencies.

**Figure 1.**
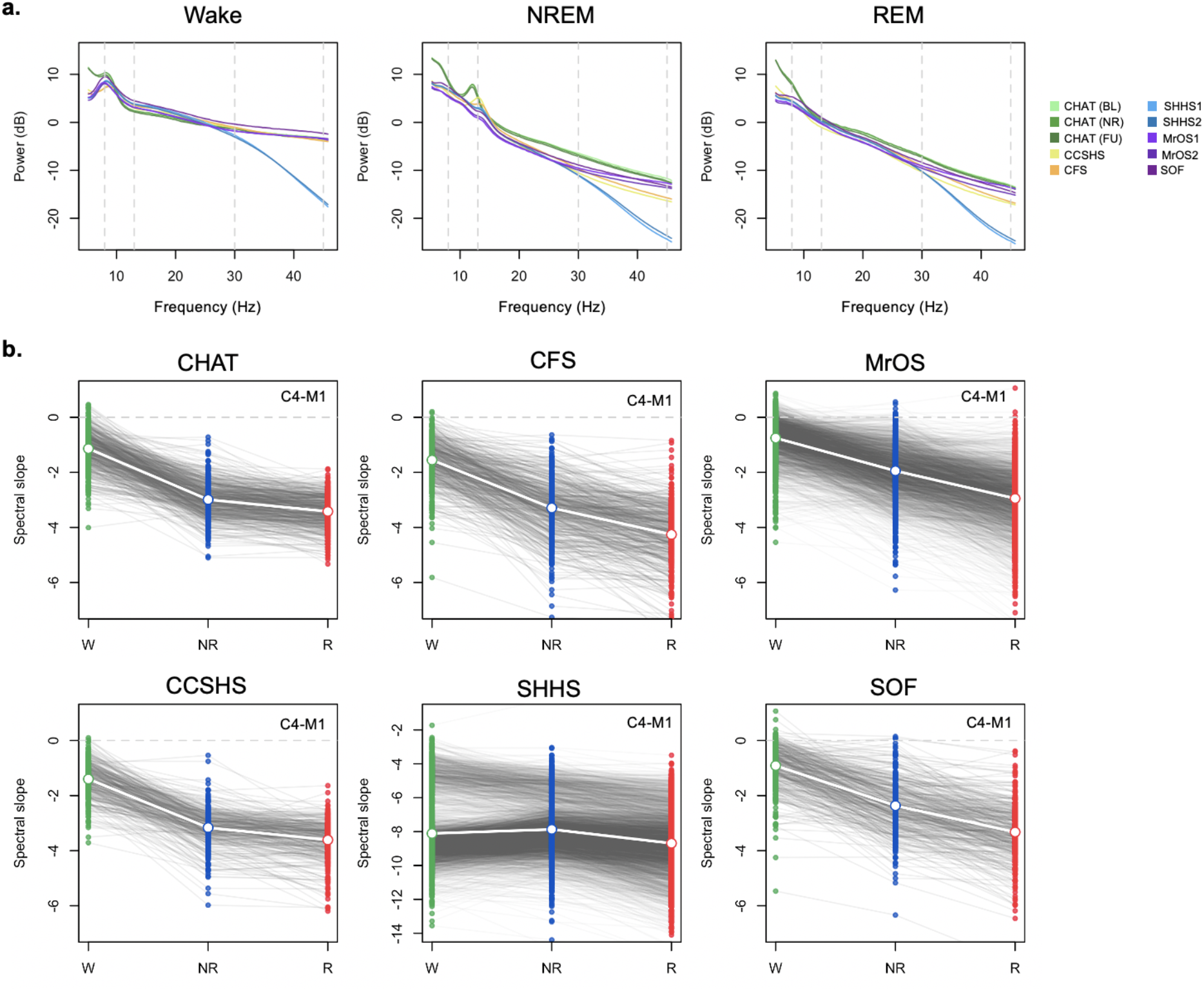
Spectral power and slopes based on contralateral mastoid (CM) referencing. For the channel C4-M1 (contralateral mastoid referencing), a) mean log power spectra (5-46 Hz) by sleep state (wake, NREM and REM) and cohort. Dashed vertical lines at 8 and 13 Hz indicate typical modes of oscillatory activity during wake (alpha rhythms) and N2 (spindles); dashed lines at 30 and 45 Hz indicate the interval within which the spectral slope was estimated. b) Estimated spectral slopes for the set of independent first-wave individuals (CHAT baseline and non-randomized samples pooled). Gray lines connect the three values for each individual. Note the different scaling of the y-axis for SHHS1 versus the five other datasets. Green, blue and red indicate wake, NREM and REM respectively. See **Table S1** for sample size details.

Evident in the raw power spectra, all cohorts other than the SHHS showed steeper 30-45 Hz slopes in NREM and REM than wake. **Figure 1b** shows estimated state-specific slopes for the six baseline cohorts. Briefly, given a power spectrum *PSD*(*f*) ~ *1/f*^*α*^, the spectral exponent *β* = *-α* was estimated as the linear slope of the log-log regression of power on frequency (see **Methods**). All three pairwise within-individual differences between wake, NREM and REM slopes were highly significant (*p* < 10^−15^ matched-pair *t*-tests) and similar results were obtained for C3-M2 (**Figure S2, Table S2**).

As implied by the spectra in **Figure 1a**, slopes were markedly steeper in SHHS compared to other cohorts (note the different *y*-axis scaling). Furthermore, most evidently during wake, the SHHS showed a bimodal distribution of slopes (also addressed in **Supplementary Information Section 1**). Given these technical issues, we elected to exclude SHHS from our primary analyses.

### Potential non-neural confounders of the spectral slope

The spectral slope in the gamma range can be affected by muscle activity as the scalp EEG is inevitably sensitive to frontalis (peak frequency between 20-30 Hz) and temporalis (40-80 Hz) ^13,22^ as well as extraocular muscles (30-120 Hz with peak at 64 Hz) executing saccadic eye movements ^23,24^. In a previous study comparing resting state EEG with and without paralysis induced by complete neuromuscular blockade, power above 20 Hz was attenuated 10- to 200-fold under paralysis, suggesting that most scalp EEG above 20 Hz is of EMG origin^25^. It also has been reported that during wake, resting state EEG recordings contaminated by muscle activity expressed flatter slopes than EEG with no EMG interference ^26^. Neither are intracranial recordings necessarily completely free of muscle activity interference ^27–30^. There are well-known differences in muscle tone and ocular movements between wake, NREM and REM sleep; muscle atonia typical to REM sleep was reported to affect frontalis muscle to the same extent as submental muscle - a standard site of PSG EMG recording^31^ but there is also evidence of increased facial muscle contractions due to limbic activation during REM^32^.

In the five cohorts excluding SHHS (CHAT, CCSHS, CFS, MrOS and SOF) we therefore investigated possible sources of bias and/or noise, first estimating state-specific slopes from 30-45 Hz chin EMG spectra. Briefly, we observed multiple, potentially non-trivial linkages between state-dependent differences in EEG and EMG slopes: 1) similar wake > NREM > REM mean differences (**Table S3**) and age-related flattening (**Figure S3**), 2) positive correlations in EEG and EMG slopes within state (**Table S4**), 3) mean differences in EMG-EEG coherence (taken to index potential contamination) between states, primarily REM > NREM > wake (**Figure S4a**), 4) significant associations between EMG-EEG coherence and EEG slope (**Figure S4b**) and 5) modest but significant associations with BMI for both EEG and EMG NREM slopes (**Table S5, Figure S5**). In the CFS, we additionally estimated 30-45 Hz slopes from the ECG, finding largely similar patterns with respect to EEG slopes as for the EMG (**Figures S6 & S7**). See **Supplementary Information Section 2** for details.

Similar to Lendner et al., conditioning on EMG slope was not sufficient to fully explain the observed, within-individual state-dependent differences in EEG slope - although we note that peaks in Lendner’s EMG power spectra at exactly 20 and 40 Hz suggest that their EMG slope estimates were themselves subject to bias/noise (see Lendner et al. Figure 2, supplement 2). Nonetheless, sources of between-individual, within-state variation (including noise and bias) may be quantitatively and qualitatively different from the sources of within-individual, between-state variation, meaning that non-neural confounds could still bias (either attenuating, or spuriously inducing) individual differences in the spectral slope.

**Figure 2.**
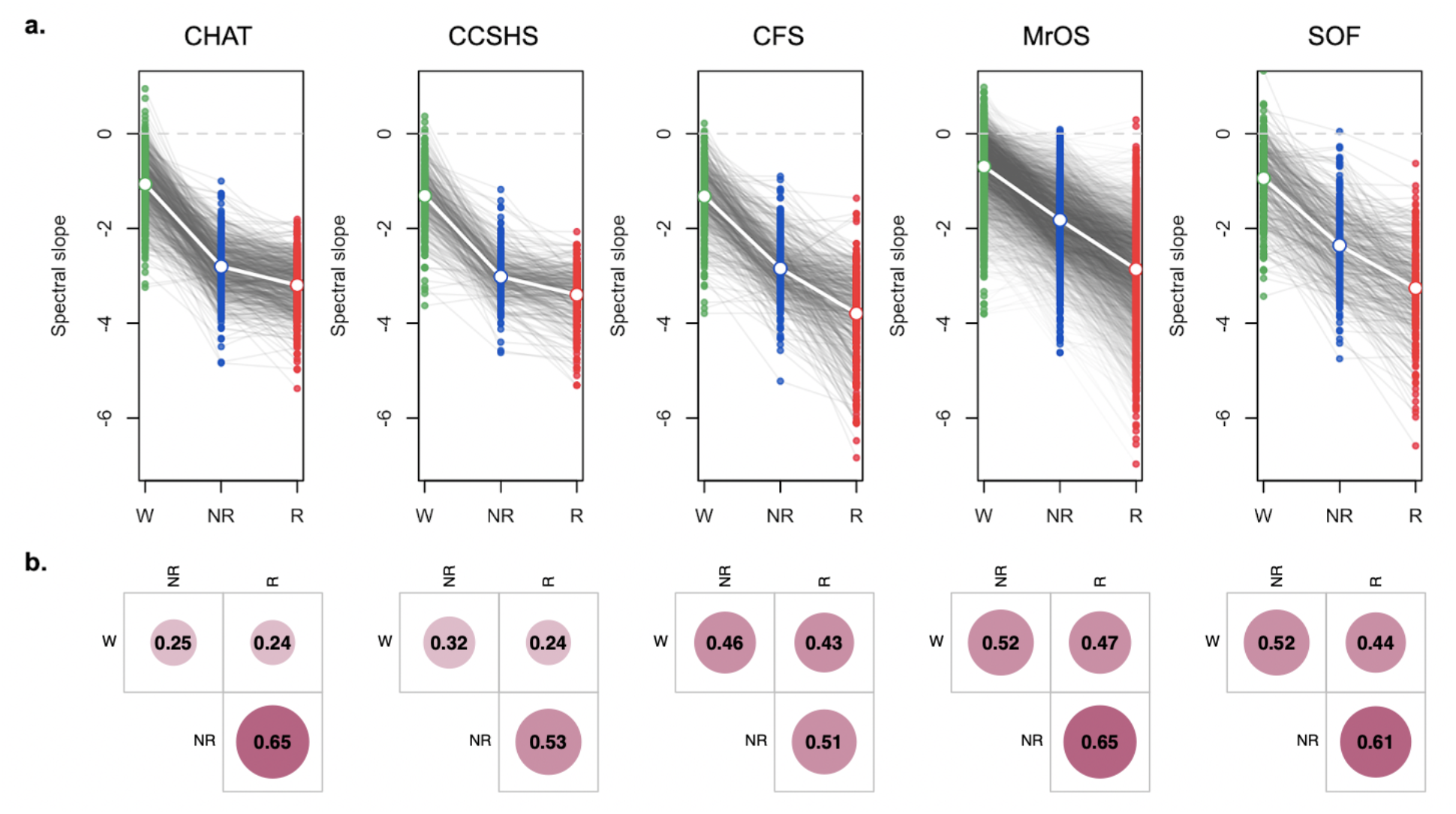
Spectral power based on linked mastoid referencing, excluding SHHS. a) Plots as for **Figure 1b**, but based on the LM referencing scheme (all state differences, matched pair *t*-test *p* < 10^**-15**^). b) Pearson correlation coefficients in slope between the three sleep states considered (all *p* < 10^**-5**^). Green, blue and red indicate wake, NREM and REM respectively.

Lendner et al. considered alternate referencing schemes including bipolar/local referencing: in our limited montage context, this corresponded to only the bipolar C3-C4 derivation. As a control, we also considered the “cross-mastoid” derivation M1-M2: although mastoid electrodes are not truly independent of neural activity, we expected any signatures of cortical arousal to be greatly attenuated, compared to the derivations involving a central scalp electrode. However, M1-M2 slopes often had greater effect sizes than C3-C4 slopes with respect to state differences (**Table S2**) and, within state, were very highly correlated with EMG slopes (**Table S4**) and BMI (**Table S5**, **Figure S5**). Given the potential role of mastoids introducing non-neural sources that might bias slope estimates, we alternatively considered a linked (i.e. averaged) mastoid (LM) referencing scheme. Although LM-derived slopes were correlated highly with contralateral (CM) slopes (e.g. for C3 *r* = 0.93, 0.88 and 0.90 for wake, NREM and REM respectively) and exhibited comparable state-related differences (**Table S8**), LM referencing (as originally employed by Lendner et al.) dramatically reduced the above mentioned markers of potential EMG/ECG-driven bias in EEG spectral slopes, e.g. as indexed by 1) EEG and EMG/ECG coherence (**Figure S6**), 2) correlation between EEG and EMG/ECG spectral slopes (**Table S7**), and 3) likely spurious associations with BMI (**Table S6**, **Figure S5**). See **Supplemental Information, Section 2** for more details.

### Linked mastoid analyses of the spectral slope by sleep state

Given concerns over a) technical factors in the SHHS and b) differences between LM- and CM-derived slopes, we based our primary analyses of the EEG spectral slope on LM referencing, with the SHHS dataset removed. We re-ran all outlier exclusions on this new dataset, yielding a final QC+ dataset of *N* = 4,459 recordings from *N* = 3,543 distinct individuals, of whom *N* = 690 (primarily in MrOS) had a second PSG.

We observed unambiguous support for statistically different slope means, namely wake > NREM > REM (**Figure 2, Figure S8 & Table S8**). Averaging over cohort means, overall *β* = −1.11, −2.58 and −3.3 for wake, NREM and REM respectively. (N1 and N3 exhibited similar slopes to N2 (−2.58), albeit typically marginally less steep, mean *β* = −2.4 and −2.34 respectively, **Table S8**).

Despite mean differences, slopes were significantly correlated across states (**Table S9**), suggesting systematic and state-independent factors influenced slope, other than arousal level *per se.* To a first approximation, N1, N2 and N3 slopes correlated r~0.7; for REM and NREM, r~0.5-0.7; for wake and sleep, r~0.2-0.5. As noted, given the broad similarity of N1, N2 and N3 slopes, all NREM analyses below used N2 sleep only, to ensure greater homogeneity in NREM sleep across individuals.

### Within-individual epoch-level discrimination of sleep state based on the spectral slope

Lendner et al. evaluated the extent to which the EEG spectral slope could be used to classify epochs as wake, REM or NREM (N3). In comparison to slow oscillation (SO) power, the spectral slope enhanced discrimination of wake versus REM, and was comparable for wake versus NREM. Here, we adopted a similar linear discriminant analysis (LDA) approach to classify epoch-level data in the CFS cohort (chosen because it contained the most diverse age range).

Given state definitions and scoring rules, it is not clear why one would expect SO power to be a particularly strong predictor of wake versus REM, however. We therefore focused on what we considered a more relevant comparison (for the question of discriminating REM from wake): different parameterizations of the higher frequency EEG, namely beta (15-30 Hz) and gamma (30-45 Hz) band power, as either an absolute or relative (with respect to total 0.5-50 Hz power) metric.

Using LDA to discriminate wake and REM in the CFS cohort, the spectral slope did not perform differently compared to beta power (*p* = 0.78, with an average accuracy of 87.5% versus 87.3% for the slope, **Figure 3**) and performed only marginally better than gamma power (*p* = 0.048, with 86.7%). In contrast, while the spectral slope performed better than beta power for classifying REM versus NREM (*p* = 10^−15^, 73.7% versus 66.8%), it was worse for classifying wake versus NREM (*p* < 10^−43^, 75.6% versus 87.2%). Although higher frequency EEG activity may - as others have suggested^33^ - be an informative (and often overlooked) feature for distinguishing REM from wake, potentially driven by the EMG content of the high frequency EEG, we did not find evidence that the spectral slope *per se* is an optimal parameterization for this particular goal. Indeed, here beta or gamma band power alone performed similarly (equivalent results were observed for relative power).

**Figure 3.**
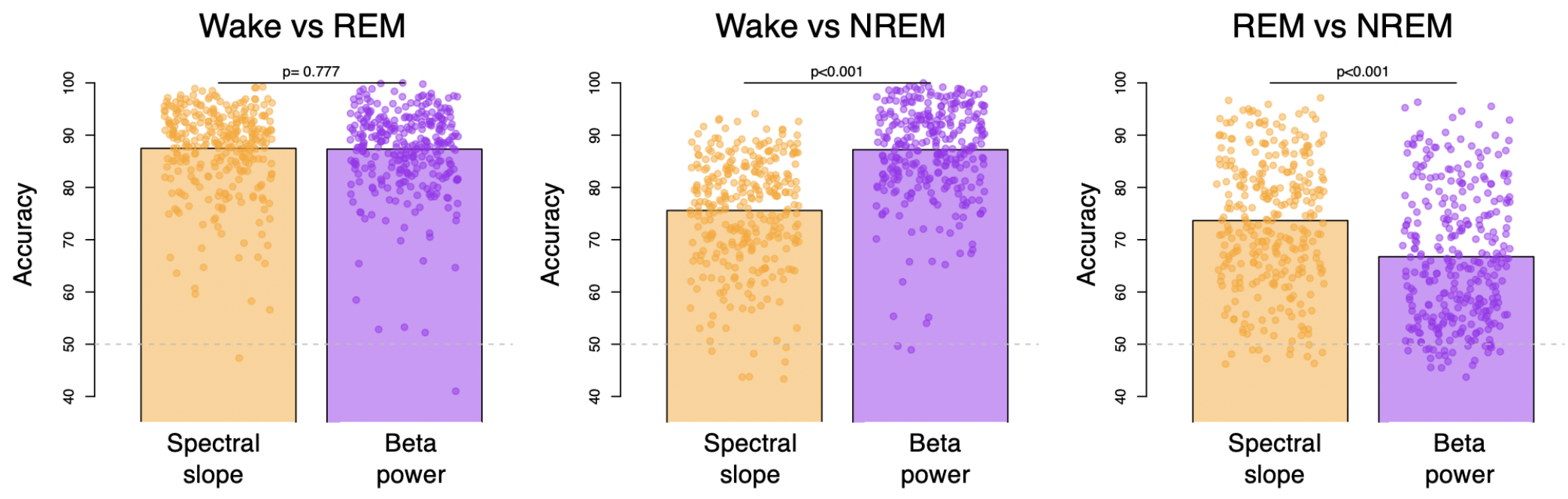
State classification using LDA based on spectral slope or beta band power. Three bar plots illustrate mean accuracies across individuals of W vs R, Wvs N2 and R vs N2 classification and dots represent individual accuracies (orange - spectral slope as a predictor and purple beta power as a predictor). P-values above the bars indicate whether there was a significant difference between accuracies produced by LDA based on spectral slope vs beta power. Dashed grey line illustrated chance level performance. See **Methods** for details.

### Variability in spectral slope by sleep state

In addition to between-state differences in means, we also characterized state-dependent differences in the variability of spectral power and slope, considering both within-individual (epoch-to-epoch) and between-individual (person-to-person differences in means) sources of variation. Analogous to **Figure 1**, but based on LM-referencing and excluding SHHS, **Figure 4** shows mean power spectra (top row) but also the variability (standard deviation units, SD) in power, partitioned into within-individual and between-individual components (middle and lower rows respectively). Across all cohorts there was consistently greater variability in waking spectral power at higher frequencies, e.g. >20 Hz, increasing up to 45 Hz, consistent for both estimates of variability. There was also a tendency for greater variability at the points of canonical oscillatory activity for wake (i.e. ~8 Hz) and NR (i.e. ~13 Hz). In contrast, variability in REM power spectra was approximately uniform across this frequency range. (**Figure S9** shows these data plotted separately for each cohort.)

**Figure 4.**
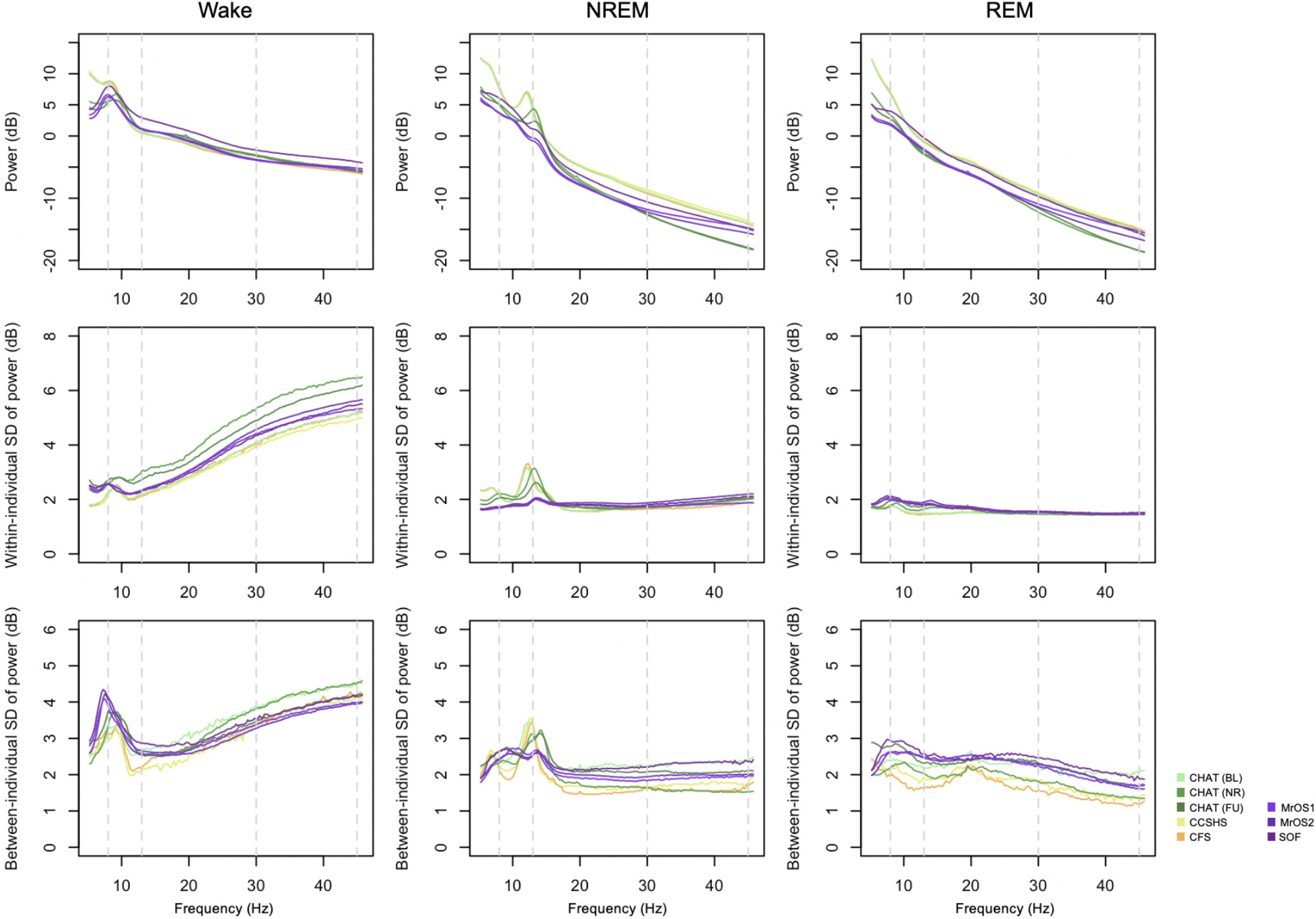
Means, within- and between-person variability in state-specific spectral power. All analyses based on the LM-referenced dataset, with the SHHS studies excluded. Within-individual variability was based on the standard deviation (SD) of epoch-to-epoch differences, calculated for each individual separately and then averaged over all individuals in each cohort. In contrast, between-individual variability was the SD based on differences between individuals’ mean power, calculated once for each cohort. See **Figure S9** for these data plotted individually for each cohort.

We next considered the components of variability in the spectral slope (**Figure 5**). Because the number of epochs observed for a given individual/state may influence the variance/error in estimated slopes (which will influence between-individual variability in mean slopes), we plotted the mean number of epochs for each state (**Figure 5**, top row). As expected, individuals typically had more NREM epochs; there was also a tendency for older cohorts to have relatively more wake than REM epochs. In general, between-individual variability in slope (**Figure 5**, second row) was similar across states, with perhaps the exception of greater variance for REM in the older cohorts (potentially reflecting their reduced REM duration). With respect to within-individual variability (**Figure 5**, third row), there was a clear pattern of greater epoch-to-epoch variability in slopes during wake compared to REM, with intermediate levels observed for NREM, which may reflect greater heterogeneity and noise in the waking data.

**Figure 5.**
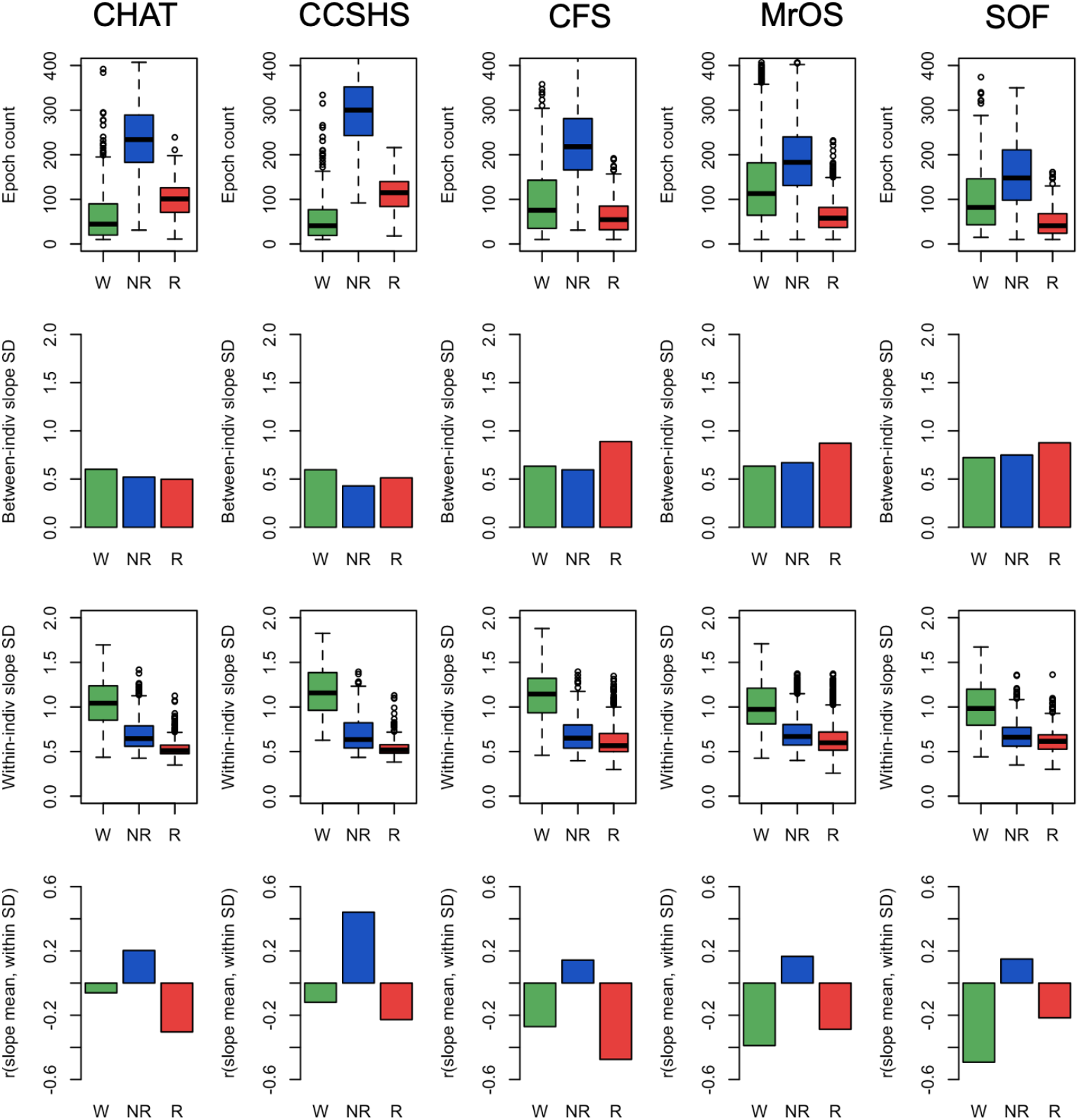
Epoch counts, variability in spectral power (within-and between-individual) and correlations between mean slope and within-individual slope SD. All analyses based on the LM-referencing; all epoch counts refer to the number of epochs passing the stringent QC procedures. Green, blue and red indicate wake, NREM and REM respectively. See **Methods** for details.

We also considered the correlation between an individual’s mean slope and the corresponding epoch-to-epoch variability in slope (for that same individual). If, for a given state/individual, epoch-level estimates of slope followed a single normal distribution, one would not expect any significant correlation between mean and variance. **Figure 5** (bottom row) shows the mean/variance correlations, averaged over individuals, separately by sleep state and cohort. Correlations were typically significantly different from 0.0, and followed a distinct pattern across cohorts: wake and REM slopes exhibited negative mean/variance correlations, whereas NREM slopes exhibited positive correlations. In other words, for REM and wake, individuals with more epoch-to-epoch variability in slope tended to have steeper (more negative) slopes, on average. The opposite was true for NREM: individuals with greater variability tended to have flatter slopes. These results suggest that looking only at a single value of the spectral slope (i.e. for one individual, an estimate based on all epochs) will miss other forms of state-dependent distributional differences in slope dynamics.

### Relationships between the 30-45 Hz spectral slope and power

The spectral slope was not independent of absolute (or relative) power across the spectrum. Consistently across all cohorts, we found that slope-power correlations showed qualitatively different patterns between wake, NREM and REM, however. **Figure 6** (top row) shows state-specific correlations between spectral slope (based on the 30-45Hz interval) and power across a broader spectrum (10-46 Hz in 0.25 Hz bins; also see **Figure S10**). For example, considering power at 30 Hz we observed highly significant (*p* < 10^−10^) negative correlations during REM, but positive correlations during wake. To visualize slope/power relationships more directly, **Figure 6** (lower three rows) also shows log-scaled absolute power stratified by a median split on spectral slope. Dashed (versus solid) lines represent mean power for individuals with steeper (versus shallower) slopes. During wake (**Figure 6**, second row), steeper slopes resulted from greater divergence in power at higher frequencies (e.g. towards 45 Hz). In contrast, during REM (**Figure 6**, bottom row) steeper slopes resulted from greater divergence at lower frequencies (10-30 Hz), which was attenuated at higher frequencies, i.e. up to 45 Hz. During NREM we observed an intermediate profile (**Figure 6**, third row). The larger differences in wake higher frequency power echoed the increased variance in power and slope (**Figures 4 & 5**). That is, individual differences in waking power and slope were driven by factors that particularly influenced gamma band power, whereas this was not the case during REM.

**Figure 6.**
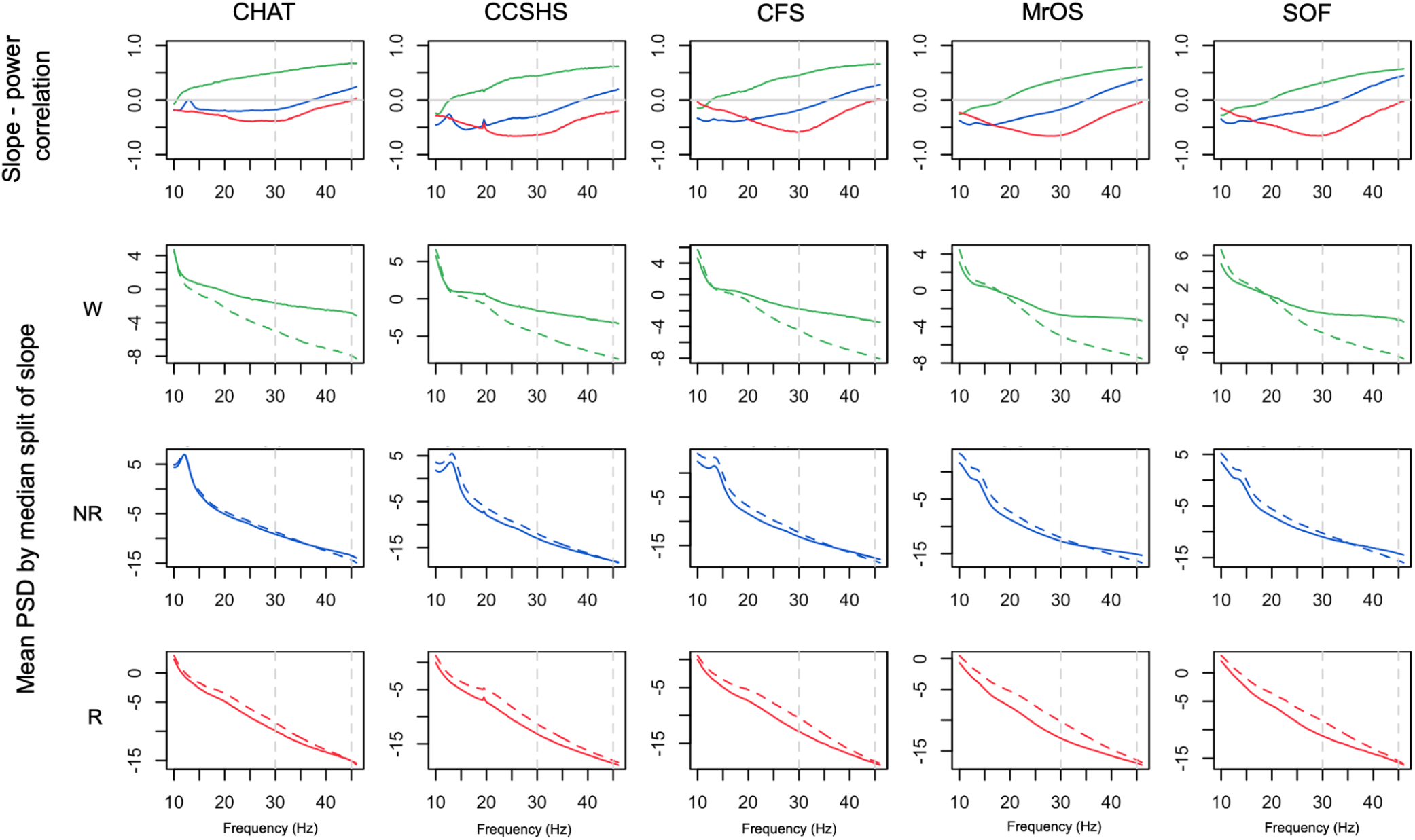
Relationships between spectral slopes and spectral power. Top row shows Peason correlation coefficients between individuals’ mean spectral power and mean spectral slope, conditional on sleep state and cohort. The lower three rows show mean power stratified by a median split on spectral power: means for the group of individuals with steeper slopes are represented by dashed (versus solid) lines. All analyses were based on the LM-referenced dataset. Green, blue and red indicate wake, NREM and REM respectively.

### Modelling state-dependent changes in the EEG power spectrum

We adopted a simplified, model-based approach to determine whether, by themselves, changes in the spectral slope (assuming a strict power law model) were sufficient to account for the (qualitatively-different) slope/power relationships depicted above, here focussing on spectral power/slope relationships (primarily **Figure 6**). We directly simulated power spectra for *N* = 5,000 individuals, initially in the form PSD(*f*) = *A/f^α^* + *e*, where *A* and *α* were normally-distributed random variables representing the spectral intercept and exponent (slope *β* = *-α*) respectively (see **Methods**). Loosely following others (e.g. refs. ^10, 11, 34^), we then extended the model in several ways; schematically depicted in **Figure 7**, these (non-mutually exclusive) parameterizations were as follows: 1) allowing spectral slope and intercept to be correlated (positively or negatively), 2) including a flat spectral component *C,* such that PSD(*f*) = *A/f^α^* + *C,* 3) allowing an alternate center of rotation *f_r_* such that log PSD(*f*) = log *A* + *α* log(*f* / *f_r_*), and/or 4) allowing for the slope to vary across the power spectrum. The latter was implemented by modelling two independent slopes (*α* and *α*\*) along with a frequency-dependent sigmoidal weight function *w*(*f*) (see the first column of **Figure S11**), such that the spectrum was in the form log PSD(*f*) = log *A* + *w*(*f*) *α* log(*f* / *f*) + (1-*w*(*f*) *α*\* log(*f* / *f_r_*). Although the second slope *α\** could in principle be parameterized differently from *α* (i.e. in terms *μ, σ, f_r_* and correlation with the intercept, in order to model a ‘knee’ in the mean spectral slope), here these were fixed to the same values for *α*. The relevant point is simply that, under these ‘tapered’ models, changes in *α* will tend to influence the slope only at higher frequencies (e.g. >30 Hz) but not at lower frequencies (e.g. <20 Hz), as modelled by *w*(*f*).

**Figure 7.**
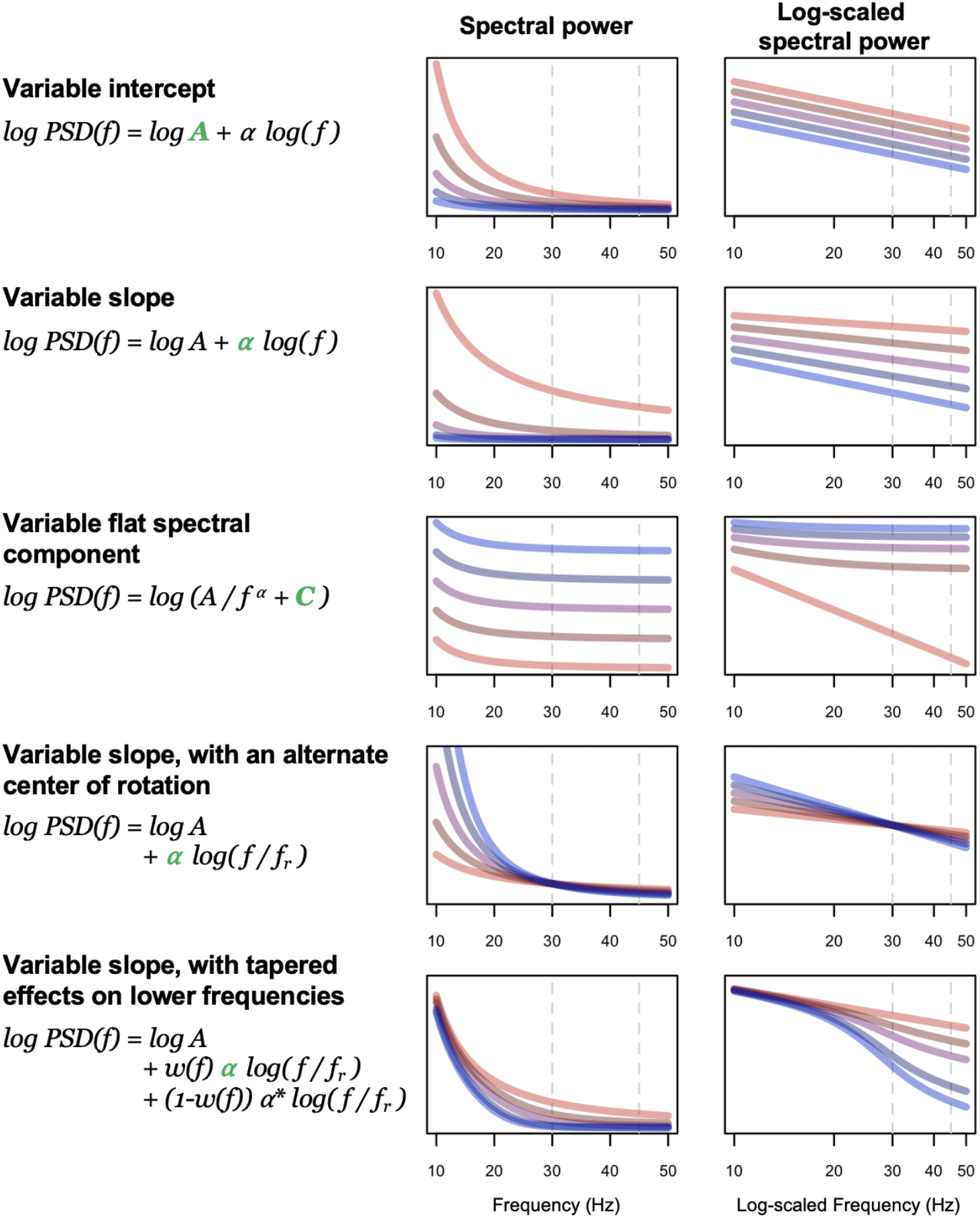
Model parameters. These cartoons illustrate the parameterizations of the aperiodic component of the power spectrum we considered in the simulations. In each case, the green term indicates the aspect of the model that was varied, and the corresponding plots show the impact of power spectra (in linear-linear and log-log coordinates, left and right figures respectively). For illustrative purposes only, the five lines (from blue to red) show the expected power spectrum for five different parameter values (e.g. α = 1, 1.5, 2, 2.5 and 3). Power absolute values/units (y-axes) are arbitrary and so not shown: these figures are intended only to show some of the qualitative patterns of differences that can arise due to variation in a given model parameter. The *w*(*f*) function was similar to those depicted in **Figure S11b**, with a 50% value at 25 Hz in this example (lower row). Beyond these factors, the model also allowed slope and intercept to be correlated, and specified a stochastic error term (smoothed with respect to frequency). See the text and **Methods** for details.

We initially set population slope parameters *α* ~ *N*(*μ, σ*^2^) to *μ* = 1, 2.5 and 3 to reflect typical observed means for wake, NREM and REM respectively. In the simulated power spectra, we followed the analysis procedures as above, estimating state-dependent variability in power, the mean slope, the correlation between slope and power, and mean power stratified by a median split on slope (**Figure 8** & **Figure S11a**). Given the consistency across studies evident in **Figure 6**, here we reproduce only the CFS results in **Figure 8** (first column), to reflect the empirical, observed values for key statistics, focussing on the state-dependent slope-power correlation and slope-stratified mean power.

**Figure 8.**
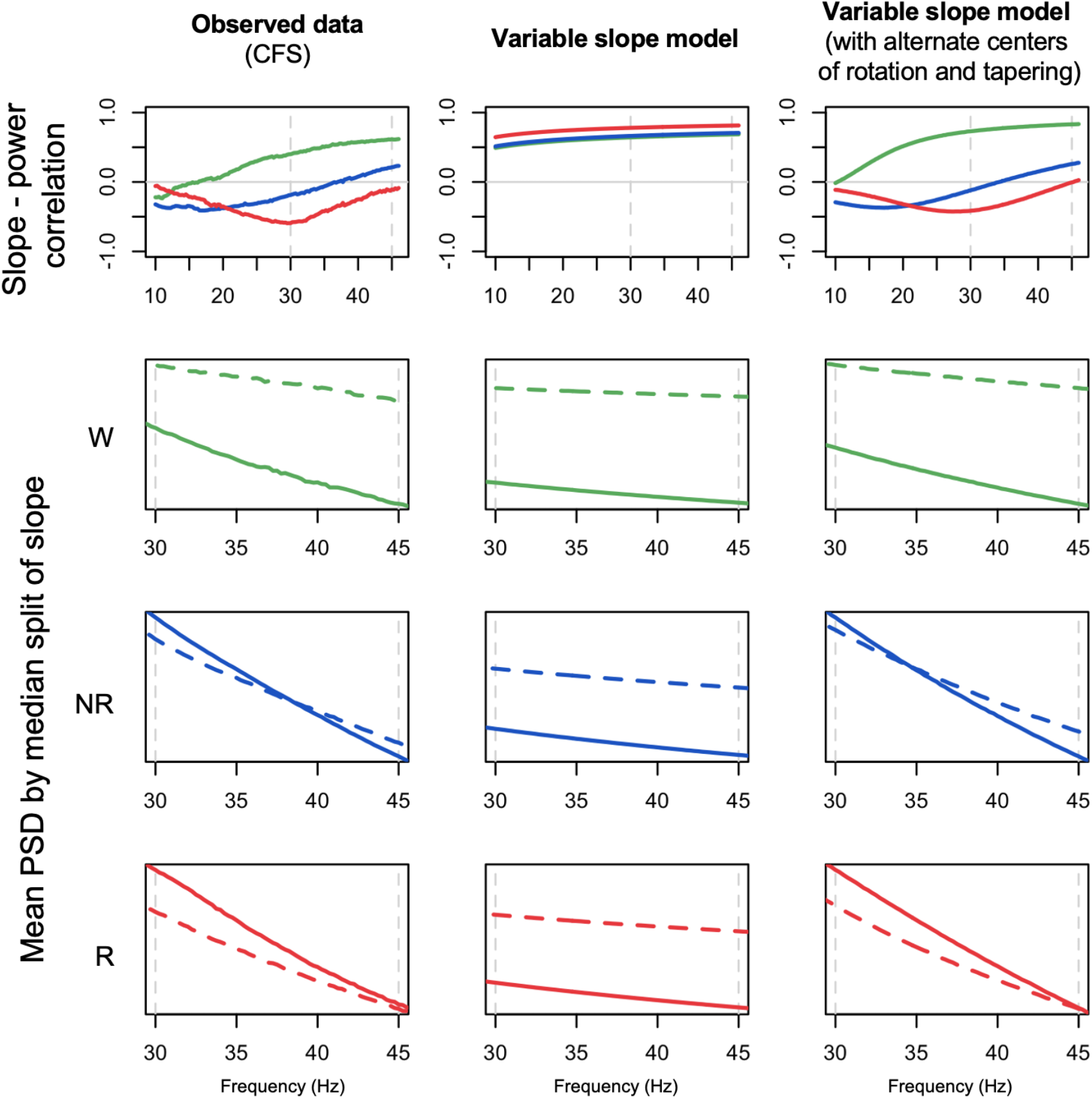
Observed data, initial and revised model simulation-based predictions. The left column of plots reproduces the observed results from the CFS cohort, for state-dependent slope-power correlations (top row) and mean power stratified by a median-split on slope (lower three rows). Green, blue and red indicate wake, NREM and REM respectively. Based on N = 5,000 simulated spectra, the middle and right columns show the equivalent simulation-based results, from a) the original model parameterization (‘variable slope model’), assuming a strict power law model with mean α = 1, 2.5 and 3 for wake, NREM and REM respectively (and SDs of 0.5, 0.5 and 0.75, approximately following the observed between-individual estimates from **Figure 5**), and b) a revised model (‘variable slope model with alternate centers of rotation and tapering’), with similar population parameters for slope means and variances but allowing different centers of rotation (*f_r_* = 10, 35 and 45 Hz for wake, NREM and REM respectively) and setting *w*(*f*) such that variation in *α* had less influence on the slope at lower frequencies (see **Figure S11b**). Whereas the initial model (simply varying mean spectral slope by state) could not recapitulate the observed results, the revised model could. See text and **Methods** for further details.

Our initial model - that assumed a strict power law implying a rotation of the slope around 1 Hz as a function of *α* (**Figure 8**, second column, and **Figure S11a**) - was unable to recapitulate the qualitatively different patterns of slope-power correlation we observed across states. Namely, slope and power were always positively correlated, meaning that individuals with steeper slopes tended to have lower power at all frequencies above 1 Hz. In a revised model, we observed two alterations that were sufficient to account for the major, state-specific pattern of results (**Figure 8,** third column): changing the center of rotation (*f_r_*) and allowing for a frequency-dependent tapering of the influence the *α* parameter on the spectral slope, via a non-uniform *w*(*f*). Specifically, we set *f_r_* to 10, 35 and 45 Hz for wake, NREM and REM respectively. Changing the center of rotation allowed for the qualitatively different slope-power relationships observed across states. However, by itself, changing only *f_r_* yielded slope-power correlations that grew very negative at lower frequencies for NREM and REM. To achieve the attenuated correlations we observed (i.e. trending back towards *r* = 0 by ~10 Hz), it was sufficient to assume *w*(*f*) functions as shown in the first column of **Figure S11b**, implying that influences on the slope at these lower frequencies were independent of factors impacting the 30-45 Hz slope.

Fully exploring these parameterizations is beyond the scope of this report and other modifications of the basic model may yield predictions that are equally consistent with our observations. These analyses were intended only to provide qualitative insights rather than quantitative fits to the data; these models also did not consider within-individual patterns of variability as documented above, parameterizing only the mean slope and spectra for each individual. Further, these models only considered the aperiodic component of the power spectrum. In real data, oscillatory activity during wake and NREM will impact observed slope-power correlations at lower frequencies (e.g. in **Figure S10**, the childhood cohorts shows dips in absolute slope-power correlations near alpha and sigma frequencies for wake and NREM respectively, presumably reflecting individual differences in alpha/spindle rhythms not strongly related to the spectral slope). Nonetheless, these analyses strongly suggest that variation in the spectral slope is not, by itself, sufficient to account for the patterns of results we observed.

### Relationships between the 30-45 Hz spectral slope and coherence

Given previous reports of reduced gamma coherence during REM^35^, as an exploratory analysis in the CFS cohort, we estimated inter-hemispheric magnitude squared coherence (between C3 and C4 using LM referencing). Absolute coherence values were generally high (reflecting the common reference) but for beta and gamma frequencies we observed significantly lower coherence during REM compared to wake, with NREM showing an intermediate pattern (**Figure 9a**). Whereas the spectral slope can be interpreted to reflect the degree of local synchronous neural activity, coherence reflects long-range functional connectivity (as well as spurious connectivity due to volume conduction). Coherence values were not independent of spectral power (here averaged across C3-LM and C4-LM), although we observed qualitatively different relationships between states (**Figure 9b**). NREM exhibited a peak in power/coherence correlation in the sigma range (presumably driven by spindle activity), but also increased coherence/power correlation above 30 Hz. In contrast, during REM sleep there was an inflection point at 30 Hz, after which individual differences in coherence and power decoupled. **Figure 9c** reproduces the slope/power correlation for CFS (as shown in **Figure 6**, but here based on the slope and power averaged over the two central channels). Finally, **Figure 9d** shows the correlations between average slope (30-45 Hz) and coherence: as for slope and power, there were qualitatively different patterns between all three states. During REM, individuals with steeper slopes tended to show higher C3-C4 coherence, particularly around 30 Hz. In contrast, during NREM, individuals with steeper slopes tended to show lower coherence at higher (>20 Hz) frequencies, whereas for wake, individuals with steeper slopes tended to show lower coherence at slower (<20 Hz) frequencies.

**Figure 9.**
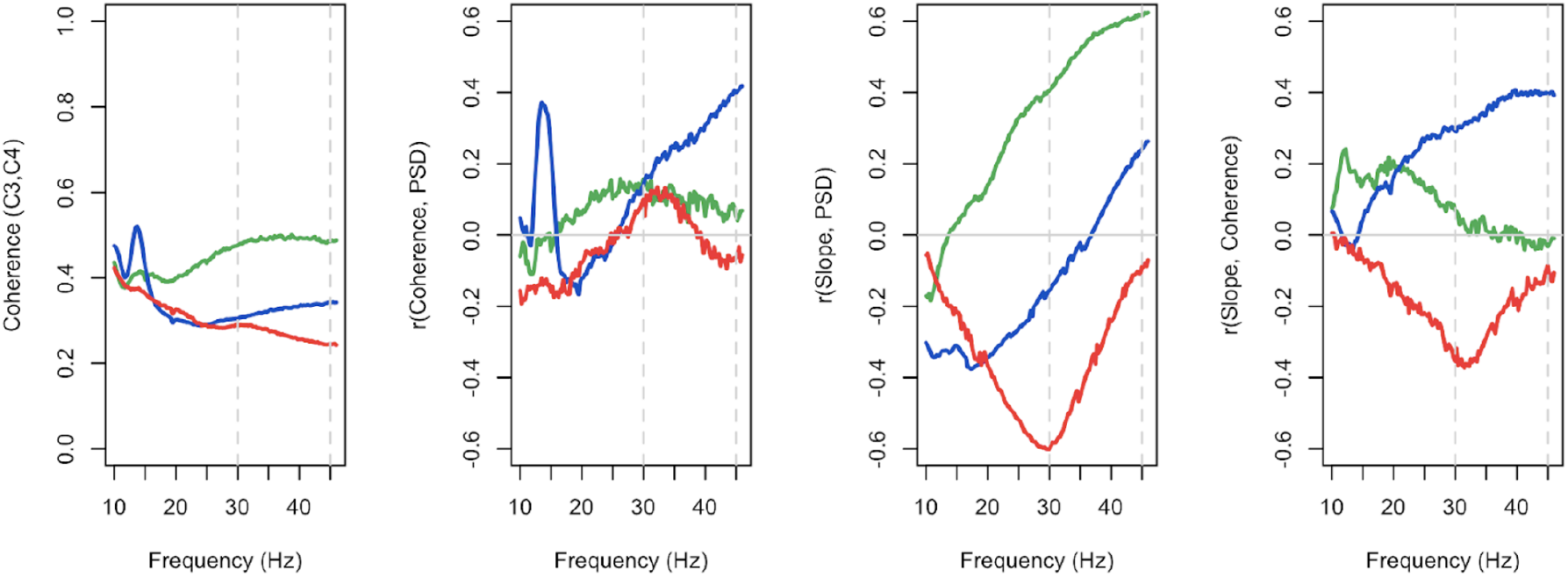
state-specific relationships between inter-hemispheric coherence, power and spectral slope. Analyses based on CFS data only, all analyses based on the LM-referenced dataset. Green, blue and red indicate wake, NREM and REM respectively. Coherence was estimated using magnitude squared coherence, see **Methods** for details.

Fully unpacking these coherence results is beyond the scope of this report (and would in any case be better executed in the context of studies with higher electrode density). Nonetheless, we point to these results - alongside the prior results for the spectral power - to underscore the types of qualitative state-dependent differences in measures related to the spectral slope, which appear to extend beyond simply differences in means.

### Demographic sources of variation in state-specific spectral slope

Finally, we considered how the spectral slope varied with age and sex, and whether those effects varied between wake, NREM and REM. First, we leveraged the repeated PSGs from the MrOS cohort of older men (~5 years between visits) to estimate test-retest reliability as well as age-related changes in a longitudinal/within-individual context. Spectral slopes showed moderate to high test-retest correlations (**Figure 10, Table S10**). For example, for C4-LM, the test-retest *r* = 0.51, 0.63 and 0.75 for wake, NREM and REM respectively (all *p* < 10^−15^, **Table S10**). We further observed significant flattening of slopes over time, albeit only for NREM and REM, with differences of 0.00, 0.32 and 0.57 for wake, NREM and REM respectively (matched pairs *t*-test *p* = 0.98 for wake, and *p* < 10^−15^ for NREM and REM). Similar patterns held in MrOS for C3-LM (**Table S10**).

**Figure 10.**
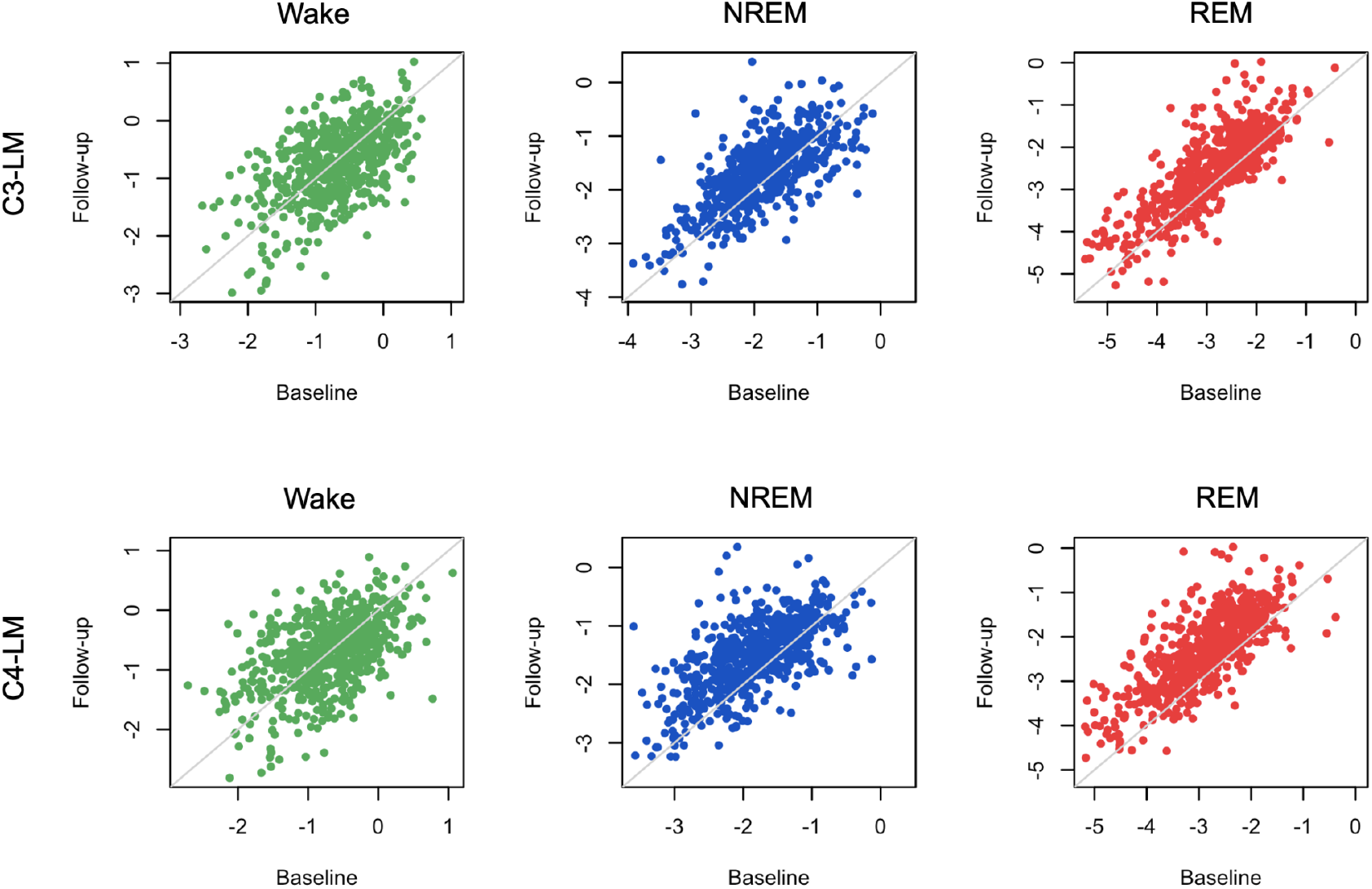
Repeated spectral slope assessment in the MrOS cohorts (waves 1 and 2). All analyses were based on the LM-referenced dataset. *N* = 610 individuals had QC+ recordings for all states in both MrOS1 and MrOS2. Visits were typically ~5 years apart (mean ages of 76.4 and 81.1 for wave 1 and 2 respectively). Green, blue and red indicate wake, NREM and REM respectively.

Furthermore, all pairwise between-state differences in slope (i.e. R-W, R-NR and NR-W) had similarly high test-retest correlations (e.g. *r* = 0.67 for R-NR, **Table S10**). In absolute terms, the magnitude of state differences grew smaller over time (all *p* < 10^−15^), meaning that states with initially steeper slopes showed greater age-related decline (i.e. REM > NREM > wake). As noted, MrOS is an elderly cohort (mean ages were 76.4 and 81.1 years in waves 1 and 2 respectively). Potentially due to either increased noise in estimated slopes during wake, or a form of floor/ceiling effect in age-related change, we did not observe significant flattening of the wake slope in this cohort. In contrast, however, REM slopes showed the greatest age-related flattening, potentially suggesting that the sleep-based spectral slope is a more sensitive measure of aging in this population.

**Figure S12** (and **Table S10**) shows the comparable results for the longitudinal component of the CHAT cohort. Although, due to the extended WASO inclusion criterion, the portion of CHAT passing QC in both waves was relatively small (*N* = 80 pairs), we still observed significant test-retest correlations, but only during sleep (for C4-LM, *r* = 0.08, 0.71 and 0.79 for wake, NREM and REM respectively, with *p* = 0.46 for wake and *p* < 10^−10^ for both sleep states). CHAT PSGs were only separated by ~6 months on average, and so mean differences may not necessarily be interpretable. With that said, a 6 month interval during childhood represents a greater developmental window, and previously we did find significant within-individual differences in spindle activity in CHAT consistent with broader cross-sectional trends seen in larger datasets ^21^. In the present longitudinal CHAT analysis, we observed nominally significant (p<0.01) age-related change, but only for a modest flattening of REM slopes (**Table S10**).

We also evaluated age-related changes in the spectral slope cross-sectionally, separately for each cohort (taking only the first recording for individuals with a repeated PSG). Results broadly pointed to age-related flattening of slopes across, and particularly for REM, although there was some degree of ambiguity and potentially inconsistency across studies. Both CHAT cohorts (pre-pubertal children, ~6-10 years) cross-sectionally showed strong age-related flattening for NREM and REM slopes, with a weaker flattening of the wake slopes (**Table S11**). Cross-sectionally, MrOS showed a flattening only of the REM slope (**Table S11**); this stronger REM effect was consistent with the longitudinal MrOS analysis above (**Table S10**). In contrast, the smaller SOF cohort of women of very advanced age (75 - 95 years) did not show any age-related changes (**Table S11**), whereas the CFS cohort (which has the broadest age range, from 7-89 years) showed an inconsistent pattern, with a flattening of NREM slope (*b* = 0.005 slope units per year, *p* = 0.006), but a steepening of REM slope (*b* = −0.01, *p* = 10^−4^) and no change in wake slope (*p* = 0.68). Secondary analyses pointed to possible non-linear age-related change for REM slopes in CFS (e.g. *p* = 5×10^−6^ for a second order, orthogonal polynomial age term, in a regression of REM slope controlling for sex, BMI, race and AHI/AI). Fuller exploration of possible non-linear age-related trends (incorporating potential cohort-specific effects also, given that these NSRR cohorts do not overlap greatly in age ranges) is beyond the scope of the current manuscript.

Finally, we observed consistent sex differences in spectral slopes, whereby males tended to have flatter slopes than females (**Table S6**, **Figure S5**). In the primary analyses, we pooled MrOS (all males) and SOF (all females) as a single cohort to facilitate the analyses of sex differences whilst controlling for cohort effects as best as possible. In the combined analysis (CHAT, CCSHS, CFS and MrOS+SOF, controlling for age, race, cohort and BMI) male slopes difered (i.e. were less steep) by 0.18, 0.34 and 0.32 units for wake, NREM and REM (*p* = 3×10^−8^ for wake, and *p* < 10^−15^ for NREM and REM, **Table S6**). As illustrated in **Figure S5**, sex differences were statistically stronger for the final LM-referenced analysis compared to the original CM-referenced analysis), although sex differences were observed consistently across all channel derivations, all cohorts and all states (**Tables S5 & S6**).

## Discussion

Using a diverse collection of cohorts from the National Sleep Research Resource, we replicated Lendner et al.’s report of progressively steeper spectral slopes (going from wake, to NREM, to REM sleep), based on the 30-45 Hz EEG. We further pointed to several technical issues that had the potential to impact estimation of the spectral slope, with a focus on potential EMG and ECG contamination, and the choice of mastoid referencing scheme. Based on thousands of diverse studies across multiple settings and with different equipment, our core results - the qualitative pattern of within-individual between-state differences in average slope - appeared to be robust to these factors.

The 30-45 Hz spectral slope is an appealing metric: as well as being easy to compute, it explicitly focuses on two components of the sleep EEG that, historically, have often been ignored: the aperiodic component, and “high” frequency activity (in our context of a typical PSG, here meaning >30 Hz). With regard to the latter point, following AASM guidelines for staging, many analyses of the sleep EEG begin by bandpass filtering in the 0.3 to 35 Hz frequency range. This is often warranted: the most easily recognisable (and oscillatory) features of the sleep EEG are <20 Hz, and it is well documented that the lower amplitude, higher frequency EEG is more susceptible to artifact ^22,36^. Nonetheless, as others have noted, even if - or precisely because - the high frequency EEG contains muscle information, it may still be informative for staging, and particularly for discriminating wake versus REM. While our study does not directly address the question as to whether including high frequency EEG improved automated staging in our cohorts, we did not find any evidence to suggest that the spectral slope *per se* was an optimal parameterization for state discrimination, compared to other simple summaries such as band power. That is, the spectral slope may remain a conceptually distinct and powerful biomarker to be used in other contexts, including within-state, between-individual analysis, but its comparative utility for sleep staging has not, in our opinion, been proven yet.

Although Lendner et al. demonstrated that slopes could be reliably estimated with different referencing schemes, our study suggested that contralateral mastoid referencing was more likely to be prone to bias and/or noise from non-neural sources. Adopting a linked mastoid referencing scheme appeared to largely (but not completely) mitigate these biases, for example, as indexed by spectral coherence between EEG and EMG/ECG, or correlation between spectral slopes derived from these different modalities. Dependencies between EEG, EMG and ECG could arise due to physiologically-driven artifact: for example, the potential for muscle artifact being picked up in the high-frequency (here >30 Hz) EEG. Alternatively, sources of noise shared across sensors (e.g. electrical noise, movement) could lead to coordinated changes in (mis-)estimated slopes. Beyond these factors, however, it remains a possibility that non-spurious, physiological linkages lead to partially coordinated slopes. There are well characterized differences between states in neural, cardiac and muscle activity, and so state-dependent changes in cortical arousal could drive and/or co-occur with other central and peripheral nervous system changes.

Importantly, the sources of variation (either physiological or artefactual) that drive individual differences in spectral slopes may difer (both quantitatively and qualitatively) from those that generate within-individual differences. That is, even if changes in slope are reliable indicators of changes in arousal levels within individuals, estimated slopes are not necessarily unbiased biomarkers of individual differences in the same phenomena. For example, if body mass index had a systematic bias on the slope (e.g. via differential cardiac/muscle contamination of the EEG), this could lead to spurious interpretations of linkages between cortical arousal and BMI, even if slope were an unbiased marker of cortical arousal level within-individual. In this spirit, we therefore sought a better understanding of the sources of variability in this metric, to realise its potential as a biomarker in research or clinical contexts, especially those involving comparisons between groups, such as neuropsychiatric patient populations.

We also discovered several issues with the EEG data in the SHHS datasets, including channel and device-specific differences. Other studies that use SHHS data - collected using very early portable EEG monitoring devices - should be aware of these efects (which we will document via the NSRR website, including the device IDs associated with differences in the spectral slopes). While these issues very directly impact analyses based on the spectral slope, other forms of analysis that implicitly use high frequency information (e.g. applying machine learning to the raw time series) could presumably be susceptible to noise and/or bias for these same reasons. We have documented these issues in the attached **Supplementary Information, Section 1.**

As well as replicating the differences in the means, we reported novel differences in other aspects of the state-specific slope distributions, including its variability (both between and within individuals) and covariation with spectral power and coherence. It is noteworthy that, despite the common characterization of REM as ‘paradoxical’ sleep (i.e. brain ‘active/wake-like’ but muscles ‘inactive/asleep’), we found that REM and wake often showed the most divergent EEG metrics, with NREM sleep being an intermediate. Specifically, as well as the mean spectral slope, we observed this pattern for 1) within-individual slope variability, 2) frequency-dependent covariation between spectral power and slope, and 3) inter-hemispheric gamma coherence. With respect to the relationships between slope and power, the observed state-dependent effects could not be accounted for by only a change in the spectral slope, under a strict power law model. Using a simplified model to provide qualitative insight, the state-specific power-slope correlations we observed were consistent with 1) changes in the center of rotation of the slope and 2) a restricted influence at lower frequencies of the factors that determined 30-45 Hz slopes. Based on a cursory initial evaluation, altering the mean and/or variance of the spectral intercept, its correlation with the spectral slope, and/or the presence of a flat spectral component ^11^ were not sufficient to account for our observations, although more work to quantitatively model our data (including any individual differences, e.g. with respect to age and sex) may be warranted.

Finally, we considered individual differences in state-specific spectral slopes, in particular test-retest stability and age-related change. In longitudinal analyses based on a subset of individuals with a repeated polysomnogram (typically ~5 years later), we observed moderate to high stability of spectral slopes, particularly during sleep. This is consistent with a prior report on a smaller sample where intra-subject reliability of spectral slope (2-25 Hz) was measured on two separate resting state recordings performed on the same day ^37^. We also observed statistically significant age-related reduction (flattening) of slopes during sleep. In cross-sectional analyses, we observed broadly consistent age-related effects in most but not all cohorts, perhaps suggesting unaccounted for between-cohort sources of variability, or nonlinear age-related effects in the CFS, the cohort with the widest age range. Flatter slopes (2-24 Hz) in older adults in cross-sectional samples were previously reported based on wake recordings during task performance ^19,20^, and similar effects of age-related flattening were observed in children (~8 yo) whose slopes were steeper during rest than in adults ^38^. Our results, however, suggested that slopes during REM appear to be particularly sensitive to age-related change. Further, the differences in slope between NREM and REM (or wake and REM) showed significant test-retest and age-related changes. Future work might investigate potential linkages between REM spectral slopes and biological aging, including the emergence of synucleinopathies including Parkinson’s disease, that are often preceded by changes in REM sleep.

Although large, this study is not without limitations. Perhaps most obviously, analyses were restricted to a very limited montage: two central channels (although Lendner et al. noted that state-related changes in the 1/*f* slope were broadly reflected across the scalp). A second caveat is that, although Lendner et al. also included wake epochs after sleep onset in their final analyses, our studies - many of which were conducted in participants’ own homes - did not systematically include any ‘quiet rest’ periods prior to sleep, and wake periods were often noisy, especially at the starts and ends of recordings. We therefore imposed a relatively strict epoch-wise filtering scheme: as such, although our results generally show a high degree of consistency, findings that point to attenuated effects during wake (namely test-retest reliability and age-related flattening) should be interpreted with this caveat in mind. Finally, the few instances of inconsistency between studies (i.e. age-related trends in the CFS cohort, which had a very broad age range and was enriched for individuals with sleep apnea) might point to factors that require different approaches (e.g. use of nonlinear modelling or more stringent QC).

Overall, as seen in other areas of electrophysiological research, theoretically-inspired alternative parameterizations of the sleep EEG have much promise, although better characterizing the sources of variation in these measures - whether from artifact, from state-related changes in arousal, or from demographically and medically relevant differences in physiology - remains an important challenge.

## Data and code availability

EEG signal analysis was performed with Luna (v0.26), an open source C/C++ package for the analysis of sleep signal data (http://zzz.bwh.harvard.edu/luna/) developed by S.M.P; all PSG data are available via the National Sleep Research Resource (http://sleepdata.org). The Luna scripts used for the primary analyses are given in **Supplementary Information Section 3**.

## Methods

### Polysomnography data

All polysomnography (PSG) data were as previously reported ^21^. Briefly, we combined PSG and demographic data on 11,630 individuals aged 4 to 97 years from the National Sleep Research Resource (NSRR). All data were collected as part of research protocols that were approved by local institutional review boards; written, informed consent was obtained from each individual before participation. The majority of individuals were from community-based samples, although two studies recruited participants for sleep apnoea. A subset (*N* = 4,079) had a second polysomnogram, typically administered 5 or 6 years after the first. We accessed European Data Format (EDF) files and annotation files (indicating manually scored sleep states in 30 second epochs as well as arousals and respiratory events). All studies used American Academy of Sleep Medicine (AASM) staging conventions, except the SHHS, which used R&K: here NREM3 and NREM4 were collapsed to a single N3 state, for consistency with the other studies.

Cohorts ordered by average age are as follows: CHAT (children), CCSHS (adolescents), CFS (wide range, but predominantly adolescents and middle-aged adults), SHHS (middle-aged adults), MrOS (elderly males) and SOF (elderly females). We generally stratified analyses by study to control for possible technical and measurement differences as well as the effects of ageing. The CHAT baseline cohort contained children with an apnea hypopnea index of 2 to 30, randomized to one of two trial arms; the CHAT non-randomized group contained children screened for the trial but not randomized, either due to unwillingness or apnea hypopnea indices that did not meet inclusion criteria. The CHAT follow-up cohort was collected 6-7 months post intervention with adenotonsillectomy or watchful waiting.

All EEG analyses were based on two central electrodes, initially re-referenced to the contralateral mastoid (C3-M2 and C4-M1). For specific analyses, we alternatively re-referenced to the average mastoid ([M1+M2]/2) denoted here as C3-LM and C4-LM (for “linked mastoid”); we also derived two bipolar channels: C3-C4 and M1-M2. The chin EMG channel was derived from left and right submentalis electrodes; the ECG channel was derived from left and right arm electrodes. All physiological signals (EEG, EMG and ECG) were resampled to 128 Hz.

### Epoch-level artifact removal

Many recordings contained extended periods of gross artifact at the beginning and ends, typically scored as ‘wake’. Although most NSRR studies did not have explicit ‘lights on/off annotations, these leading/trailing wake periods typically included ‘lights on’ periods, i.e. with the participant not in bed. To alleviate this issue of excessive artifact during wake, we removed all leading/trailing wake epochs, meaning that all ‘wake’ data reported below occurred during the sleep period (i.e. after sleep onset and before the final sleep epoch). We further removed any epoch containing a manually annotated arousal, apnoea or hypopnea. All subsequent analyses were state-dependent; we only selected epochs of a given state if they were flanked by at least one other similar epoch on both sides, to exclude transitional/unstable periods of sleep. Of note, these criteria meant that a non-trivial proportion of the overall sample was excluded, and in particular the studies of younger individuals (because of their relatively low rate of extended WASO).

We next applied epoch-wise normalization to each channel, subtracting its median value. EEG channels were further high-pass filtered at 2 Hz using a zero-phase Kaiser window FIR (transition bandwidth 2Hz, ripple 0.01). Separately for each stage (N1, N2, N3, R or W), we removed epochs for which either C3-M2 or C4-M1 had a) a flat signal spanning more than 10% of the epoch, b) a clipped signal spanning more than 10% of the epoch, c) more than 10% of the epoch exceeding 100 μV, d) more than 1% of the epoch exceeding 250 μV, or e) a maximum absolute amplitude less than 5 μV. Based on EEG and chin EMG channels, we further removed epochs that were statistical outliers (+/- 3 standard deviation (SD) units from the mean) for at least one of those channels for one or more Hjorth parameter (activity, mobility, complexity)^39^, by comparing that epoch to the mean across all epochs of that stage and channel. Hjorth-based statistical outlier removal was performed twice. Finally, we removed epochs that were flanked by multiple already-masked epochs (where the masking may have been due to any of the above reasons). Specifically, we required that at least 3 of 5 flanking epochs passed QC, either for the 5 preceding or the 5 following; we further required that retained epochs were immediately flanked by at least one other QC-passing epochs.

Collectively, this procedure was intentionally conservative, weighing specificity over sensitivity with respect to selecting clean and homogeneous epochs for the final analytic samples. Because we wanted to select a single set of criteria to be applied across all studies and stages, this means that studies had a non-trivial proportion of epochs removed (especially for younger individuals who did not meet the WASO criterion). Future analyses of these datasets focussed only on the spectral slope during sleep could of course achieve larger sample sizes by ignoring the extent of wake in each study.

### Spectral power analysis

We used the Welch method to estimate power spectra per epoch, using a 4-sec0nd segments each with 50% (2-sec0nd) overlap, applying a Tukey (cosine-tapered) window with a=0.5, yielding a spectral resolution of 0.25 Hz. We also estimated spectral band power using the following definitions: slow (0.5 - 1 Hz), delta (1-4 Hz), theta (4-8 Hz), alpha (8 - 11Hz), sigma (11-15 Hz), beta (15-30 Hz) and gamma (for this analysis, defined as 30-45 Hz to match the interval in which the spectral slope was estimated). In sensitivity analyses, we compared spectral slopes estimated from the Welch spectra to those based on multitaper analysis; here we applied 29 tapers and set the time half bandwidth product to 15 to achieve a frequency resolution of 1 Hz for a 30 second epoch. The resulting spectral slopes correlated *r* > 0.99 and so we based all final analyses on the Welch method.

As artifacts (e.g. corresponding to electrical line noise at 60 Hz and sub harmonics) can lead to sharp peaks in power spectra which may bias the estimation of spectral slopes, we quantified the extent of spectral “peakedness” within the 30-45 Hz interval. Specifically, we detrended log-transformed power spectra (assuming a linear x-axis frequency scale) and applied a median filter (using an 11 point = 2.5 Hz window) to smooth the detrended spectra. Labelling the detrended spectra *D* and the detrended and smoothed spectra *S,* we quantified peakedness as the kurtosis of the *D* - *S* difference spectra. Power spectra with strong peaks will show more leptokurtic (long tailed) distributions.

As well as summarizing mean power per individual/channel/stage by averaging over all epochs, we estimated the standard deviations of epoch-level metrics, to facilitate the analyses of within-individual variability in spectral power.

### Spectral slope estimation

Following Lendner et al., to estimate the spectral slope we extracted the absolute power from 30 to 45 Hz (here 61 values) and fit log-log linear regression models, which they showed to be appropriate for this interval of the EEG spectrum, being free of strong oscillatory activity. To reduce the impact of line noise and other artifacts that might induce sharp spectral peaks, we removed outlier data points (i.e. particular frequency/power pairs): specifically, prior to running the main regression, we fit an initial model and removed points that had residuals greater than 2 SD units from the mean. Also, because we estimated slope only over a relatively restricted interval (30-45 Hz), we did not consider the potential issue that can arise with different representations of lower versus higher frequencies due to the log-scaling of linearly uniform frequencies. For example, there is only a 1.45-fold difference in the log-scale intervals from 30 to 31 Hz compared to 44 to 45 Hz; in contrast, there is >20-fold difference between 30 to 31 Hz and 1 to 2 Hz intervals, implying that if we had looked at a broader frequency range, this issue would be more apparent.

Power spectra were estimated individually for each epoch. To derive the spectral slope, we averaged log-scaled spectra, and then calculated a single slope estimate. As a sensitivity analysis, we also calculated two other estimates: i) averaging untransformed spectra, then taking the log and estimating the slope; alternatively, ii) averaging slopes calculated per epoch on the log-transformed spectra. That is, the three approaches can be labelled: *slope(mean(log(X))), slope(log(mean(X)))* and *mean(slope(log(X)))* respectively, where *X* are the epoch-level power spectra. An advantage of the third approach is that it also provides a convenient estimate of the variability in slope across epochs. We expect qualitatively similar results from all approaches, although the second approach may be more sensitive to outlier epochs not otherwise filtered out. Work on the resting state eyes open/closed alpha rhythm differences suggested that log-transforming power at the epoch level prior to averaging was optimal in that context ^40^. Based on an initial analysis in the CFS cohort, all three metrics were effectively equivalent, both in terms of correlation with each other (*r* > 0.95) and the magnitude of tests of mean differences between stages (data not shown). The one exception was that *slope(log(mean(X)))* was less correlated with the other two metrics during wake (here *r* ~0.7), which presumably reflects a greater sensitivity to noisy/outlier epochs that occur more often during wake. As noted, our final analyses were based on *slope(mean(log(X))),* although we do not expect any substantive differences in results if these alternatives were used.

### Individual-level outlier rejection

After obtaining stage-specific slope and spectral power estimates for all individuals, we removed individuals from all primary analyses if they either 1) did not have a sufficient duration of epochs for key stages, or 2) had excessive outliers for key metrics. Specifically, we required at least 5 minutes (10 epochs) of nominally artifact-free data for W, N2 and R (as key analyses were based on these stages, and some individuals had relatively short N1 and/or N3 durations). We then removed individuals who were outliers (4 SD units) for any of the following metrics (applied sequentially such that latter filters only consider the set of currently of included individuals): gamma band power for C4-M1 or C3-M2 for W, N2 or R; gamma band power for a mastoid-mastoid derivation M1-M2 for W, N2 or R; the kurtosis (spectral peakedness) measure for C4-M1, C3-M2 or M1-M2 for W, N2 or R; the mean and SD of the spectral slope estimates for C4-M1 or C3-M2 for W, N2 or R; for C4-M1 or C3-M2, the difference in spectral slope between R and N2, between R and W or between N2 and W; finally, the mean and SD of the EMG spectral slope. For the second round of analyses based on linked mastoid channels, the C3-LM and C4-LM were additionally included in the above exclusions, as well as C3-M2 and C4-M1.

Our primary analyses are presented with these relatively stringent individual-level outlier rejection criteria; however, all key results were qualitatively similar if less stringent (or if no) outlier rejection procedures were applied instead (data not shown).

### State classification

To classify individual epochs in the CFS dataset based on the spectral slope (or other metric), we used linear discriminant analysis (LDA, as implemented in “MASS” R library) together with leave-one-out cross validation. Classification was performed in a pairwise fashion for W vs R, W vs N2 and R vs N2. Accuracy was computed as the measure of classifier performance. Individuals with less than 20 epochs for any of the states were excluded from this analysis, resulting in a final sample of 311 individuals. Following Lendner et al., to ensure that the chance level equated to 50% accuracy, we randomly downsampled one state to ensure the same number of epochs for both states. To reduce variability, we repeated the previous step 50 times, averaging to obtain the final accuracy per individual. Within individual, spectral slopes and log-scaled power estimates were z-scored prior to LDA. Matched pair *t*-tests were used to compare accuracies based on spectral slopes versus band power.

### 1/*f* model-based parameterization of individual differences in power spectra

We directly simulated power spectra for *N* = 5,000 individuals, initially in the form PSD(*f*) = *A/f^α^* + *e*, where *A* and *α* were random variables for the spectral intercept and exponent (slope *β* = *-α*) respectively. Following others ^10,11^, we also extended the basic power law equation model in three ways. First, whereas a strict power law implies the slope rotates around 1 Hz (i.e. log(1) = 0), we allowed an alternate point of rotation, *f_r_* to be specified, such that log PSD(*f*) = log *A* + *α* log(*f* / *f_r_*) + *e*. Second, following ref. ^11^ we allowed for an additional ‘flat’ spectrum *C,* as a single, fixed term added to all points of the spectrum, such that log PSD(*f*) = log(exp(log *A* + *α* log(*f* / *f*) + *e*) + *C*). Third, we allowed the magnitude of the influence of *α* to vary across the power spectrum, by weighting its contribution by a frequency-dependent sigmoid weight function, *w*(*f*) = 1/(1 + exp(-(*f* - *w_f_*) / *w_s_*)), meaning that at frequency *w_f_* the impact of the slope defined by the random variable *α* is 50% of its full value; it rises to 100% at higher frequencies, and at lower frequencies, falls to towards 0%. The scaling factor *w_s_* determined the sharpness of transition. An independent random variable *α\** was defined similarly to *α* and applied with weight 1-*w*(*f*). In this way, although the overall, marginal properties of the spectra were broadly similar, this parameterization allowed for variation in the slope in one frequency range (e.g. >20 Hz) to be independent of variation in the slope at lower frequencies.

After generating a sample of *N* = 5,000 independent spectra (0.5 to 50 Hz in 0.5 Hz increments) under a particular set of parameter values, we used log-log linear regression to estimate the realized spectral slope in the 30 - 45 Hz frequency range, for each individual spectrum. In order to replicate the results shown in **Figure 6**, we then calculated the Pearson correlation coefficient between the slope and power value (for power from 0.5 to 50 Hz) across all spectra; based on a median-split of the estimated spectral slope, we also plotted the mean power spectra for individuals with ‘steep’ versus ‘shallow’ slopes.

## Supplementary Information

### Section 1: Technical factors in the SHHS datasets

Compared to other cohorts the SHHS exhibited marked differences in mean power spectra, as well as a bimodal distribution of C4-M1 spectral slope, presumably reflecting technical issues with the recordings. Indeed, SHHS wave 1 data collection occurred 1995-1998 (wave 2 2001-2003) and relied on an early generation of amplifiers - Compumedics P-series Sleep Monitoring devices - that were more prone to artifact; for example, Compumedics noted potential issues with amplifier grounding. The P-series was not a truly digital system and, instead of being collected against a common reference, EEG inputs were hardwired into the amplifier interface and labelled EEG (C4-M1) and EEG2 (C3-M2), and so could not be re-referenced offline. Proprietary algorithms were applied to signals in a manner that was not readily transparent to SHHS study staff. Although the SHHS are described in the NSRR as having only a high-pass hardware filter set at 0.15 Hz, the power spectra in **Figure 1a** suggest these devices had a non-uniform frequency response over this range, leading to the supralinear spectral slope on the log-log scale across the 30 - 45 Hz range (with a ‘knee’ around 30 Hz). The estimates of spectral slope were highly deviant for both SHHS waves: for example, for C4-M1, the mean wake *β* values were −8.1 and −7.6 for wave 1 and 2 respectively, in contrast to values close to −1.0 for all other cohorts (as expected for wake^41^).

We also observed a second type of difference within the SHHS: although still discrepant compared to other cohorts, estimated slopes for C3-M2 were not as steep as for C4-M1 (wake *β* = −5.5 and −5.5 for wave 1 and 2 respectively, **Figure S1a**). Furthermore, the bimodality in spectral slope noted for C4-M1 was not present for C3-M2 (**Figure S1a**). In SHHS1, approximately 50 recording units were used, each for ~100 participants; after refurbishment, the same physical devices were used in the smaller SHHS2. We found that device ID had a marked impact on the spectral slope for C4-M1 but not C3-M2, accounting for the bimodal spectral slope distribution (**Figure S1b**). Most individuals would not have had the same physical unit across both studies: we noted it was the same units (rather than the same individuals) who were outliers in each wave (**Figure S1b**).

These results suggest that 1) technical factors (presumably related to device frequency response characteristics, or subsequent low-pass filtering) impacted SHHS recordings in a manner that greatly biased the 30-45 Hz spectral slope, although 2) the effect differed between (hardwired) C4-M1 and C3-M2 channels, and 3) a further layer of device-to-device variability specifically impacted C4-M1 for ~6-7 of the ~50 devices used. As well as the 30-45 Hz spectral slope, these technical factors could potentially bias any analyses that (implicitly or explicitly) rely on the high (>30 Hz) frequency content of the SHHS EEG, e.g. machine learning on raw EEG time series for automatic staging. For these reasons, as well as the inability to re-reference to linked mastoids, we excluded the SHHS from the majority of the analyses.

Nonetheless, although not a focus here, we note that the SHHS may still provide usable data for certain types of spectral slope analysis if one assumes that any technical differences were constant throughout the recording. Inasmuch as filtering reduced power by a frequency-dependent constant multiplicative factor, relative differences in log-scaled slopes may be expected to be preserved. For example, the REM - N2 *relative slope* distribution was unimodal and broadly similar across channels and SHHS waves, and also in comparison to other cohorts. Furthermore, motivated by the prior work on anesthesia, we hypothesized that benzodiazepine use (sedatives that lower brain activity) might be associated with steeper (more negative) slopes. Indeed, the 181 individuals in SHHS1 who reported regular benzodiazepine use had significantly steeper C3-M2 slopes, with similar effects across wake (*b* = −0.25, *p* = 1×10^−5^), NREM (*b* = −0.30, *p* = 2×10^−7^) and REM (*b* = −0.33, *p* = 2×10^−4^), controlling for age, sex, BMI, race, AHI and arousal index.

### Section 2: Mastoid referencing and potential sources of non-neural bias in the spectral slope

In the five cohorts excluding SHHS (CHAT, CCSHS, CFS, MrOS and SOF) we investigated possible sources of bias and/or noise, first estimating state-specific slopes for the 30-45 Hz chin EMG spectra. Across all states, although EMG slopes were flatter than EEG slopes, they exhibited the same W > NREM > REM pattern (means of −0.21, −0.53 and −0.82 respectively, all *p* < 10^−15^; also see **Table S3**) and were less steep in older cohorts (**Figure S3**). Within sleep state, EMG slopes were positively correlated with EEG slopes: for example, controlling for age, sex, race and BMI the standardized coefficients from a regression of C3-M2 slope on EMG slope were *b* = 0.04, 0.08 and 0.07 (SD units) for wake, NREM and REM respectively (all *p* < 10^−15^, see **Table S4**). Taking spectral coherence as an index of potential EMG-EEG cross-talk, we observed a state-dependent wake < NREM < REM pattern for beta band coherence across all cohorts (**Figure S4a**), and EMG-EEG beta band coherence was negatively correlated with the EEG spectral slope during sleep (**Figure S4b**), meaning that people with higher cross-talk between EMG and EEG tended to have steeper EEG slopes.

As aspects of body type (e.g. degree of excess fat) can impact EMG sensitivity and may modulate the extent of non-neural artifact in the EEG ^42^, we asked whether body mass index (BMI) was associated with spectral slope, which might indicate the influence of non-neural sources. EMG and EEG slopes both showed modest but significant associations with BMI during NREM sleep, *b* = −0.01 per BMI unit (*p* = 0.004) for C3-M2 slope, and *b* = −0.02 (*p* = 4×10^−11^) for EMG slope; similar associations were also observed in the SHHS dataset (**Table S5, Figure S5**).

We observed similar between-state patterns of mean spectral slopes (wake > NREM > REM) for C3-C4 as for C3-M2 and C4-M1, but also for M1-M2. Of note, the cross-mastoid channel often showed comparable effect sizes (and corresponding within-individual *t*-statistics) as for the original contralateral mastoids, and often greater efect sizes than for C3-C4 (**Table S2**). M1-M2 slopes were also more highly correlated with EMG slopes (**Table S4**) and showed particularly strong correlations with BMI (**Table S5**, **Figure S5**).

In the CFS cohort, we additionally estimated 30-45 Hz spectral slopes for the ECG, finding that wake slopes were less steep compared to NREM and REM slopes (matched pairs *t*-test *p* < 10^−15^, with means of −4, −5.1 and −5.2 for wake, NREM and REM respectively). ECG slopes were significantly correlated with EEG slopes, e.g. for C3-M1 *r* = 0.15, 0.20 and 0.18 for wake, NREM and REM (*p* = 0.002, 4×10^−5^ and 2×10^−5^); these ECG-EEG slope associations persisted when additionally controlling for age, sex, race, BMI, AHI and AI (data not shown).

In the CFS cohort, we estimated state-specific magnitude squared coherence between all EEG derivations and either the EMG or ECG (**Figure S6**). Particularly during sleep, there was high coherence between EMG/ECG and M1-M2, e.g. over 0.4 for EMG, and over 0.6 for ECG, with peaks in the sigma/beta band, but extending into gamma frequencies. Males tended to show higher EEG-EMG/ECG coherence than females. Whereas the C3-C4 did not show strong coherence with EMG/ECG channels, both contralateral mastoid channels (C3-M2 and C4-M1) showed relatively high coherences (up to ~0.4), presumably reflecting the extent to which EMG/ECG artifact was picked up at the mastoids.

Given the potential role of mastoids introducing non-neural sources that might bias slope estimates, we alternatively considered a linked (i.e. averaged) mastoid referencing scheme. As noted, whereas Lendner et al. originally used linked mastoid (LM) referencing, we adopted a contralateral mastoid (CM) scheme so as to be able to include SHHS data. That is, instead of C3-M2 and C4-M1 we used C3-(M1+M2)/2 and C4-(M1+M2)/2, denoted here as C3-LM and C4-LM. LM-derived slopes were correlated highly with CM-derived slopes (e.g. for C3 *r* = 0.93, 0.88 and 0.90 for wake, NREM and REM respectively). Across states, LM slopes were, on average, slightly less steep (difference of ~0.05 log power/log Hz) but the magnitudes of state-dependent differences were similar between LM- and CM-derived slopes.

**Figure S6** shows that C3-LM and C4-LM exhibited only very low EMG/ECG coherences (<0.1). Using linked mastoids eliminated the associations between NREM slope and BMI previously observed for contralateral mastoids, C3-C4 and M1-M2 (**Figure S5**, **Table S6**). Although still statistically significant, correlations between EEG and EMG slopes were reduced (to less than half the effect size during sleep) when based on linked mastoids (**Table S7**), and EEG-ECG slope correlations were eliminated (all *p* > 0.05). With regard to cardiac activity at mastoid channels due to their proximity to the carotid arteries, ECG potentials on the scalp have opposite polarities in the left and right hemispheres and are slightly asymmetric due to heart position, with higher amplitude on the left^43^. As illustrated in **Figure S7**, LM referencing takes advantage of this to effectively cancel out much of the cardiac activity seen at each mastoid (conversely, the cross-mastoid M1-M2 derivation exacerbates it).

### Section 3: Luna script used in the primary spectral slope analyses

#### Primary Luna command file

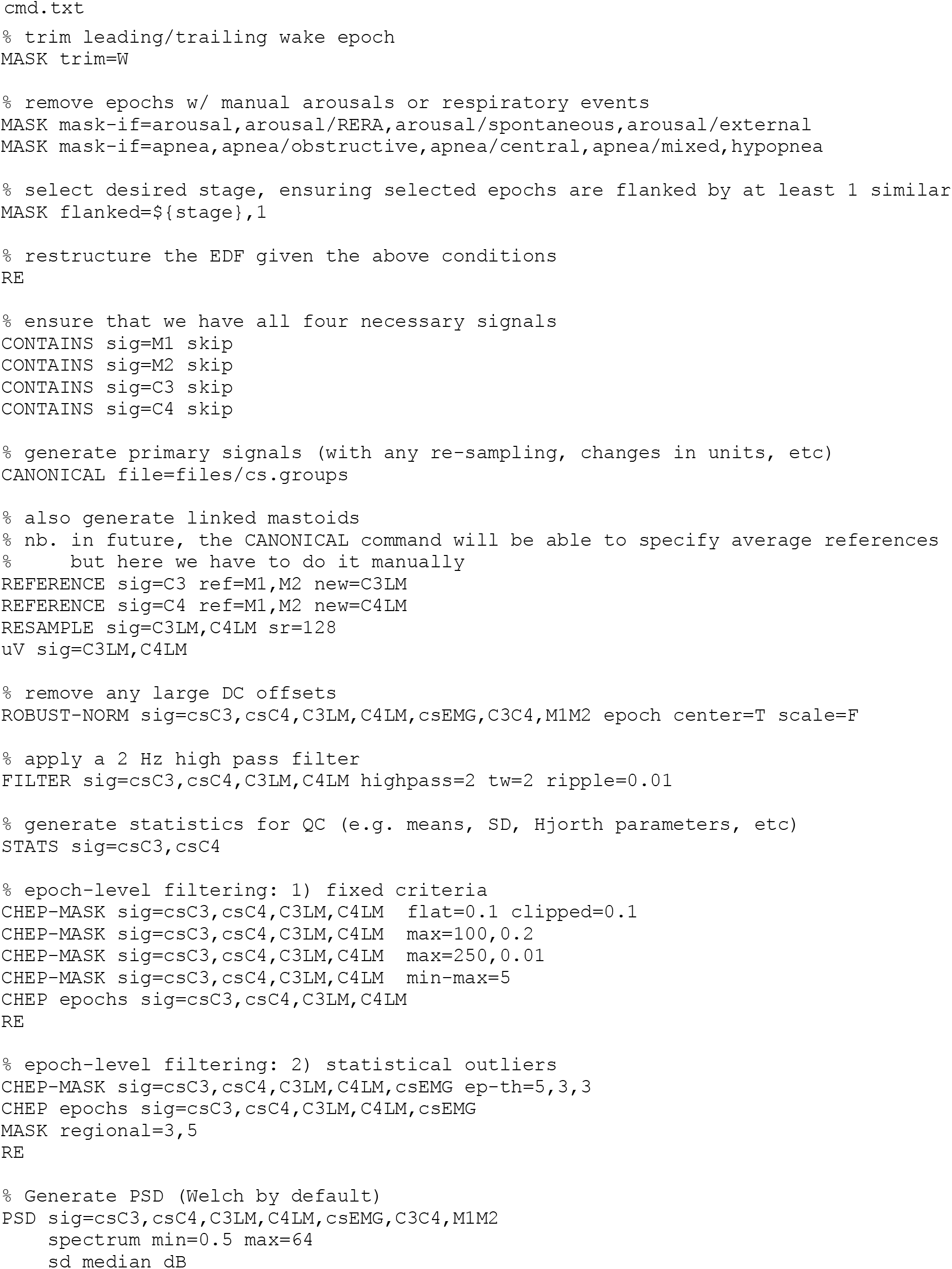
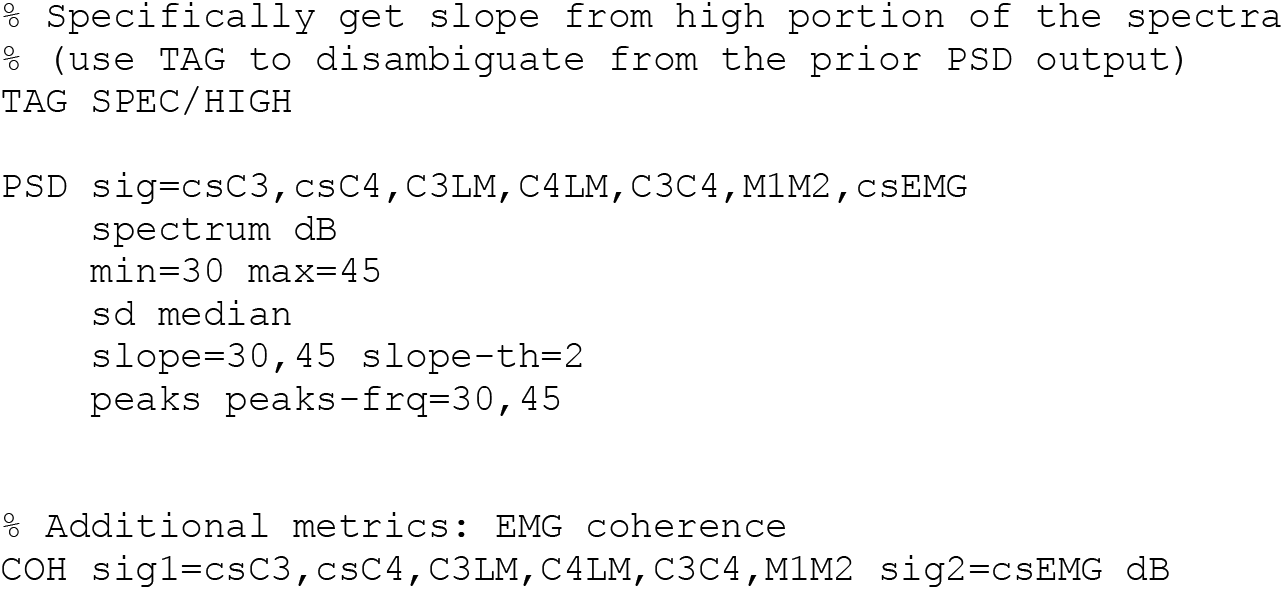

#### Canonical signal definitions

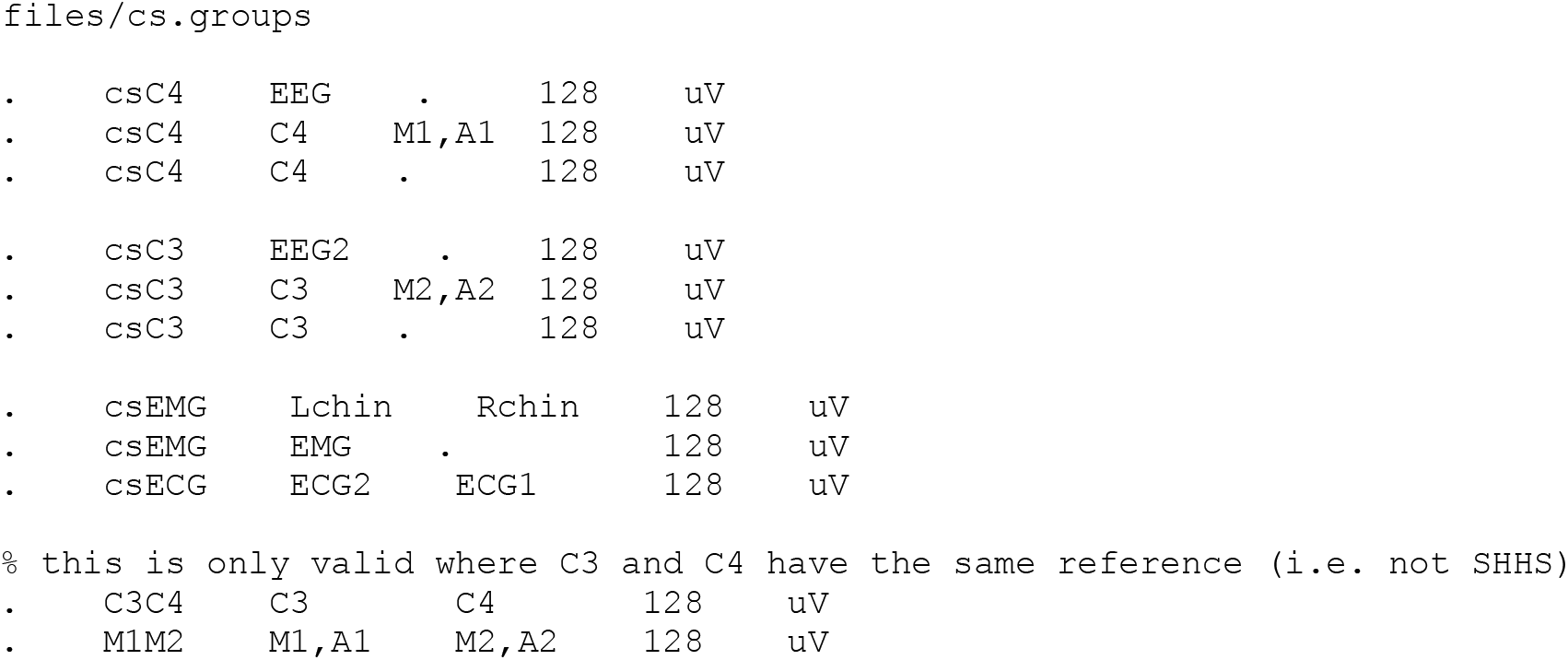

#### Invoking Luna

e.g. for N2 sleep if `s.lst` was the ‘sample list’ (IDs, EDF file and annotation file locations for one or more individuals):

~~~
luna s.lst -o out/n2.db gamma=30-45 stage=N2 < cmd.txt
~~~

(nb. in practice, jobs were parallelized)

See http://zzz.bwh.harvard.edu/luna/ for details on Luna command syntax.

### Section 4: Supplementary Figures and Tables

**Figure S1.**
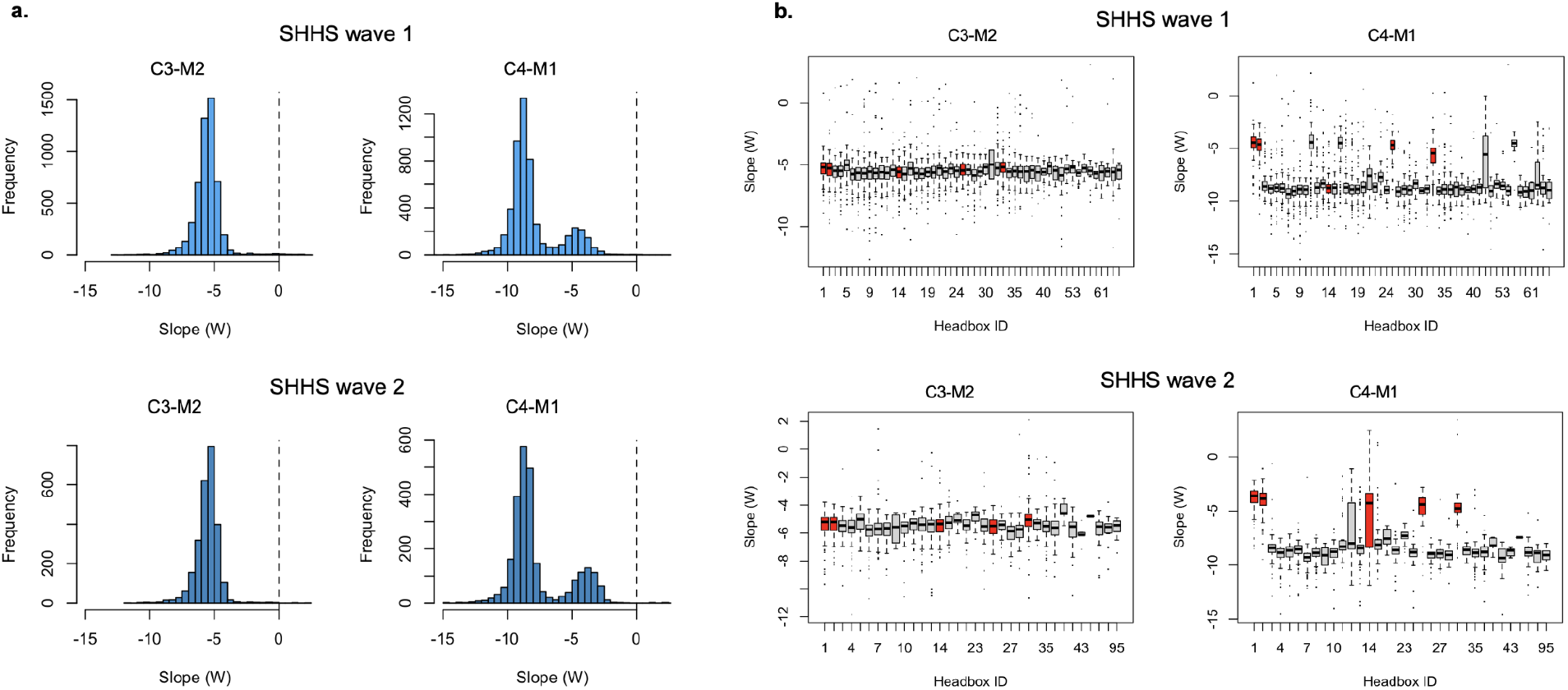
Spectral slopes in SHHS. a) Histograms of W spectral slopes for C3-M2 and C4-M1, for both wave 1 and 2, which indicate a bimodal distribution for C4-M1 only. b) Mean spectral slopes (separately for C3-M2 and C4-M1 in wave 1 and 2) stratified by the ID of the physical recording device (headbox). The same units and IDs were preserved across waves 1 and 2 (albeit with fewer individuals/devices used, as well as a handful of new devices introduced for wave 2). The devices that were outliers for C4-M1 in wave 2 were also outliers in wave 1.

**Figure S2.**
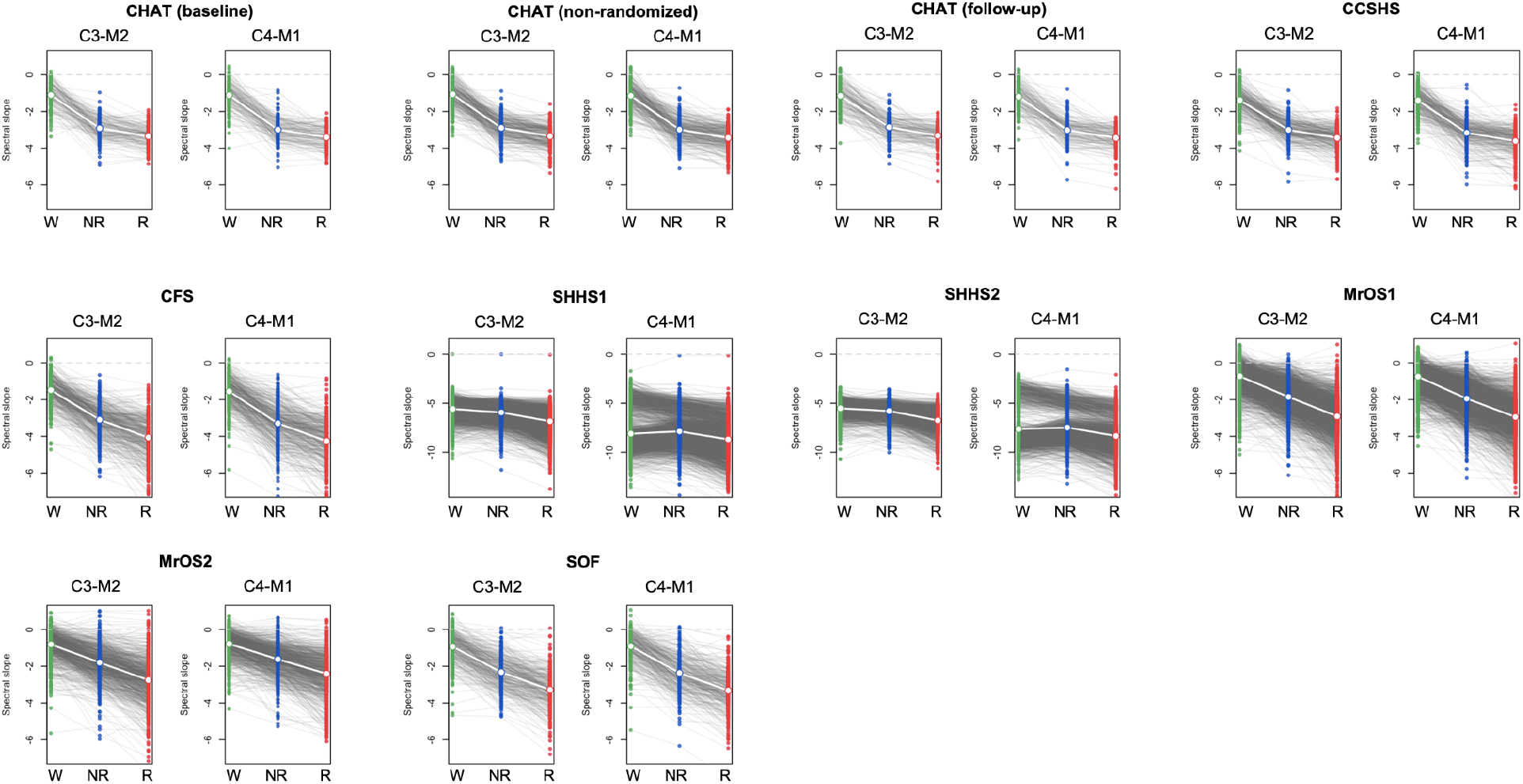
EEG spectral slopes (CM-referenced dataset). See legend for **Figure 1** for details: similar methods were applied to generate these plots, but here listed for both central channels, and separately for all ten samples. Green, blue and red indicate wake, NREM and REM respectively.

**Figure S3.**
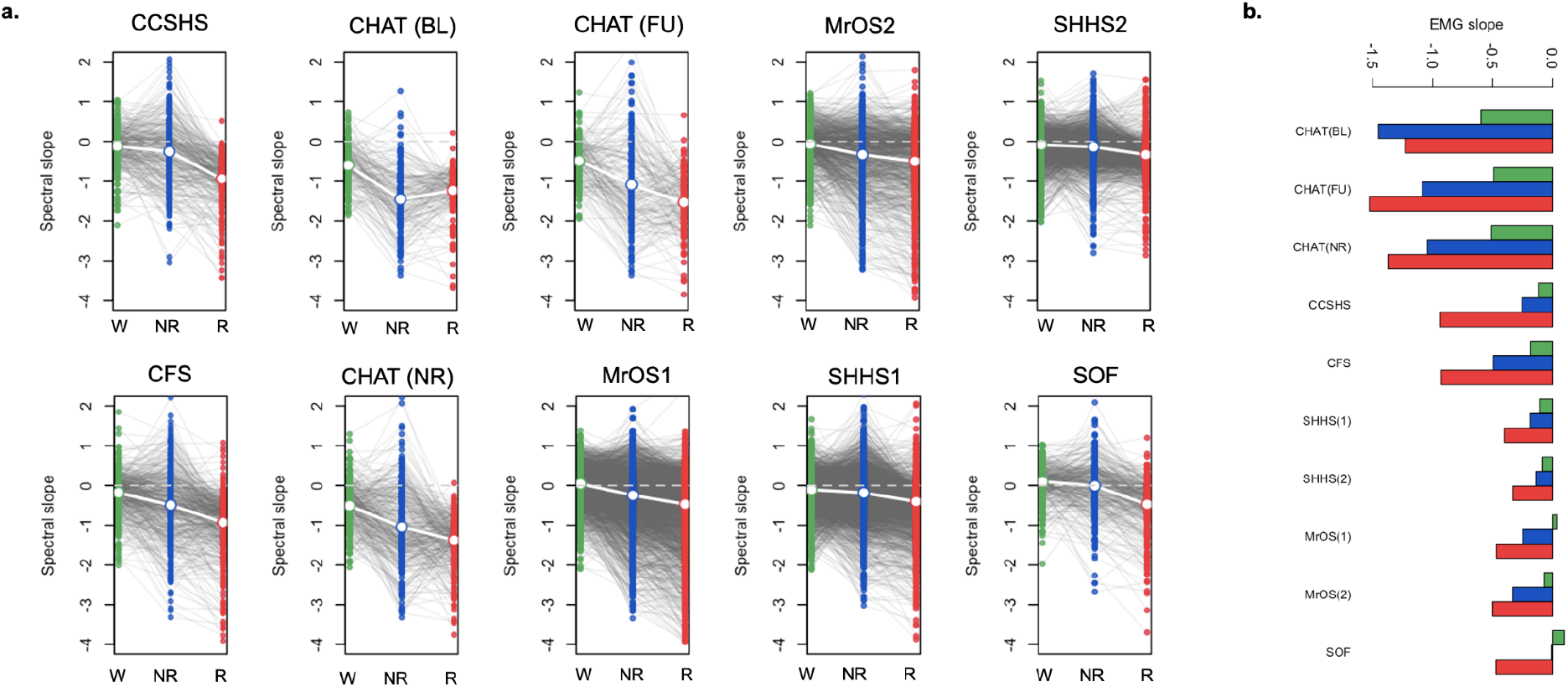
Spectral slope of the EMG. a) Distributions of EMG 30-45 Hz spectral slopes, stratified by state and cohort. b) The mean EMG slopes (identical to those in panel a.) plotted differently, to emphasize the age-related flattening. Green, blue and red indicate wake, NREM and REM respectively. Also see **Table S3**.

**Figure S4.**
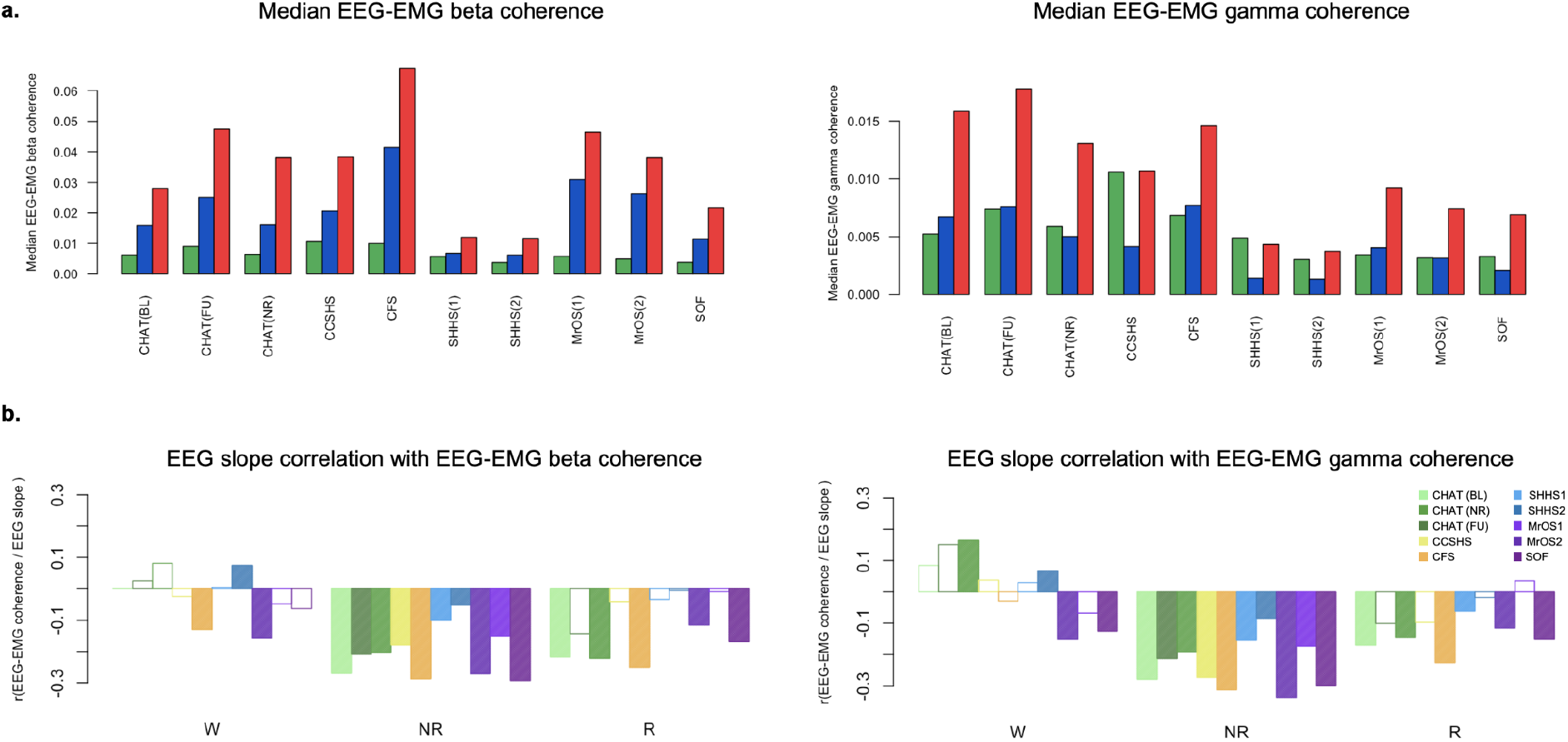
EEG-EMG coherence and EEG spectral slopes. a) For beta and gamma bands, median EEG-EMG coherence stratified by state and cohort. All analyses used CM-referencing. b) Correlations between EEG slope and EEG-EMG coherence, by state and cohort. Particularly during NREM, individuals with higher EEG-EMG crosstalk (i.e. coherence) tended to have steeper slopes. Green, blue and red indicate wake, NREM and REM respectively.

**Figure S5.**
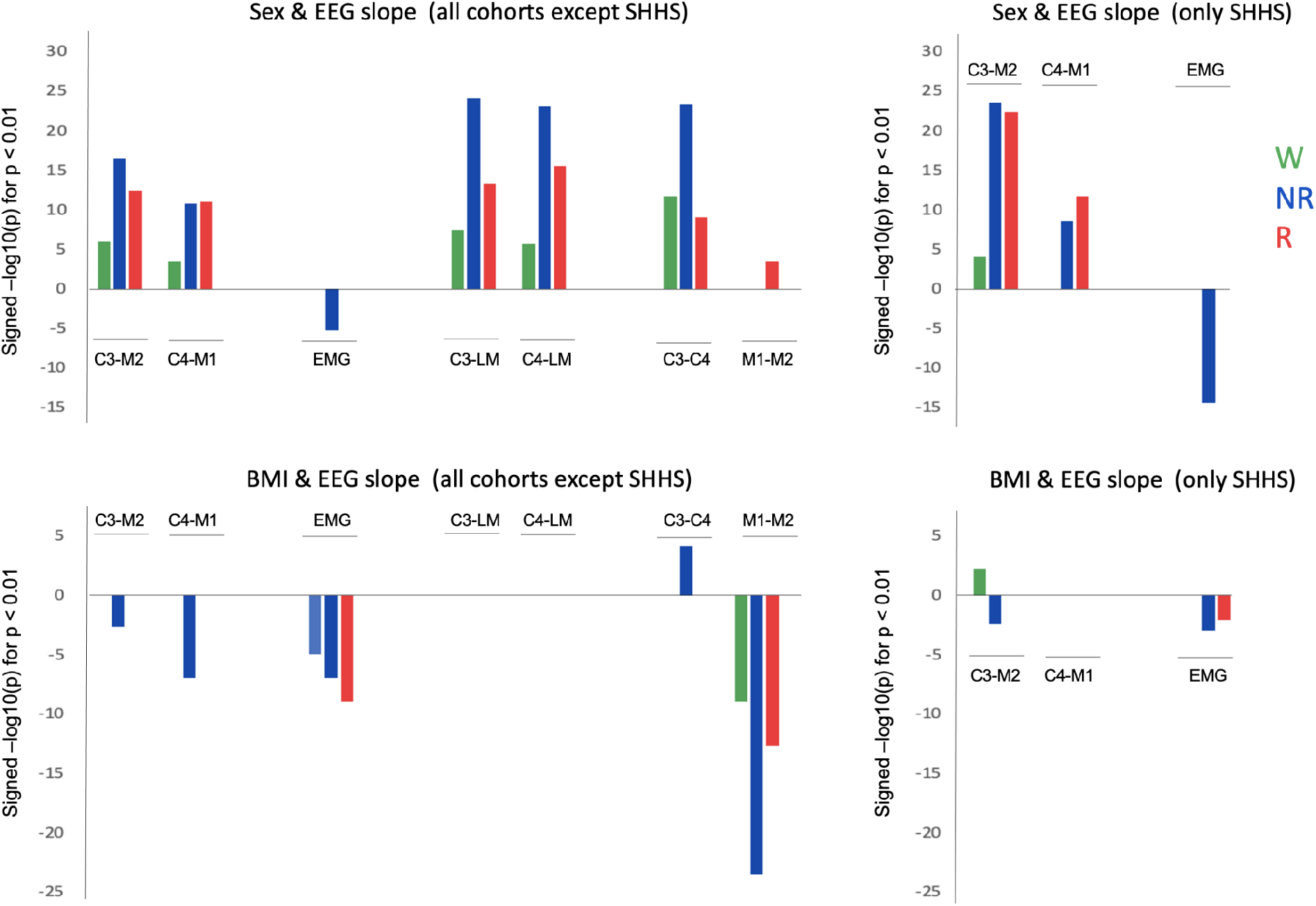
Summary of sex & BMI associations with the EEG spectral slope. Results indicate the signed −log10 *p*-value (as an index of relative effect size) from regressions of slope on sex, BMI and other covariates (see **Methods**). Only associations with *p*<0.01 are shown. Green, blue and red indicate wake, NREM and REM respectively. Also see **Tables S5** & **S6.**

**Figure S6.**
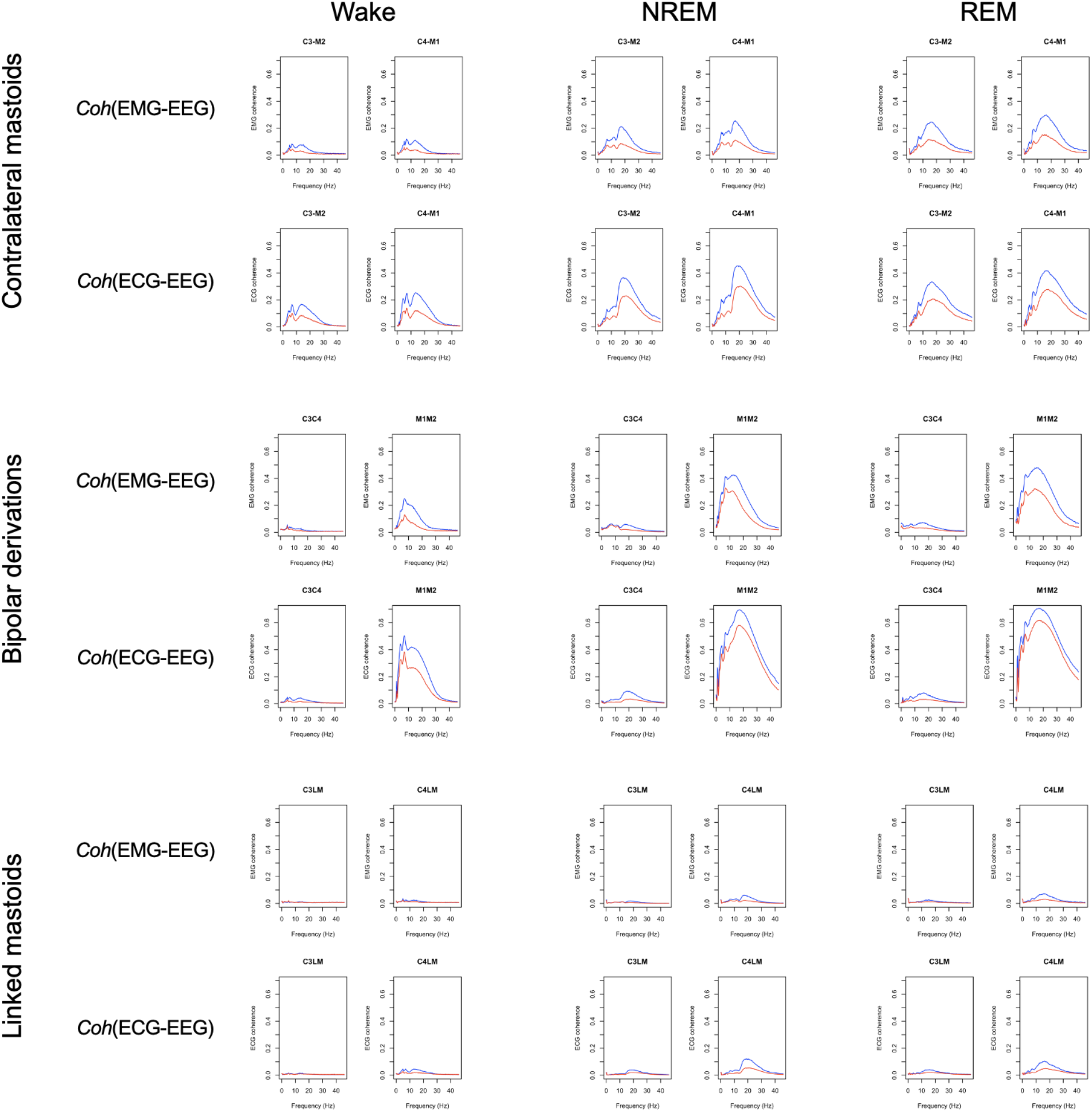
Coherence between EEG and EMG/ECG in the CFS cohort. Blue/red lines indicate mean coherence for males/females. Note that the use of linked mastoid referencing appeared to reduce EMG/ECG contamination, as indexed by spectral coherence with the EEG.

**Figure S7.**
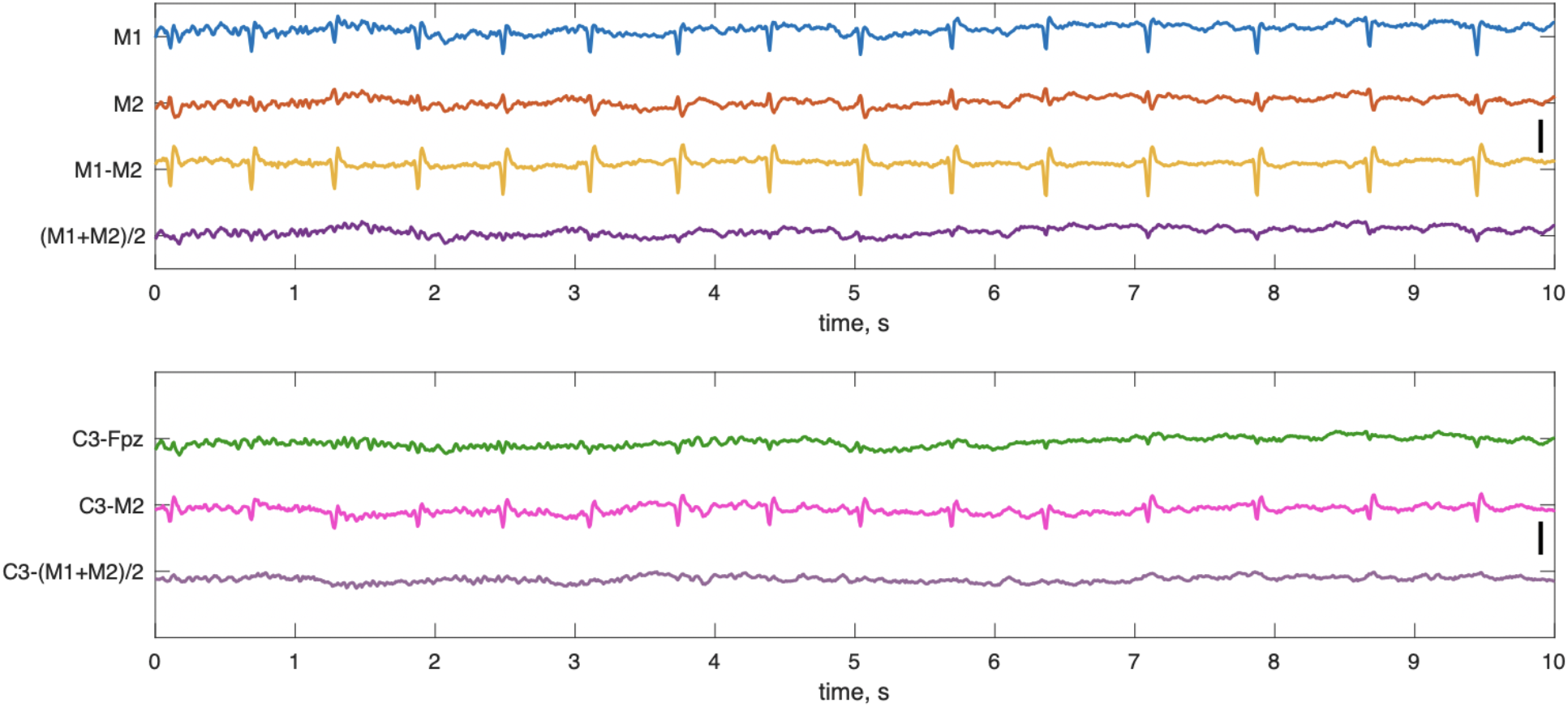
Choice of reference and ECG artifacts. The upper plot illustrates 10s of N2 from M1-Fpz, M2-Fpz, M1-M2 (the ‘cross-mastoid’) and (M1+M2)/2 (linked mastoid) for a subject from CFS with extreme cardiac interference in C3-M2. Fpz was the recording reference electrode. Both M1 and M2 time series are severely affected by ECG artifacts. Due to the opposite polarity, however, the linked mastoid (M1+M2)/2 signal almost fully eliminates this issue. The bottom plot shows the C3 channel with different referencing (the recording reference Fpz, M2 and (M1+M2)/2) during the same period. Note how the contralateral mastoid reference introduced cardiac artifacts into the C3 channel; however, if C3 in fact contained strong cardiac artifacts prior re-referencing, using the contralateral mastoid would have helped to cancel it out. The vertical black bar on the left corresponds to 100 uV.

**Figure S8.**
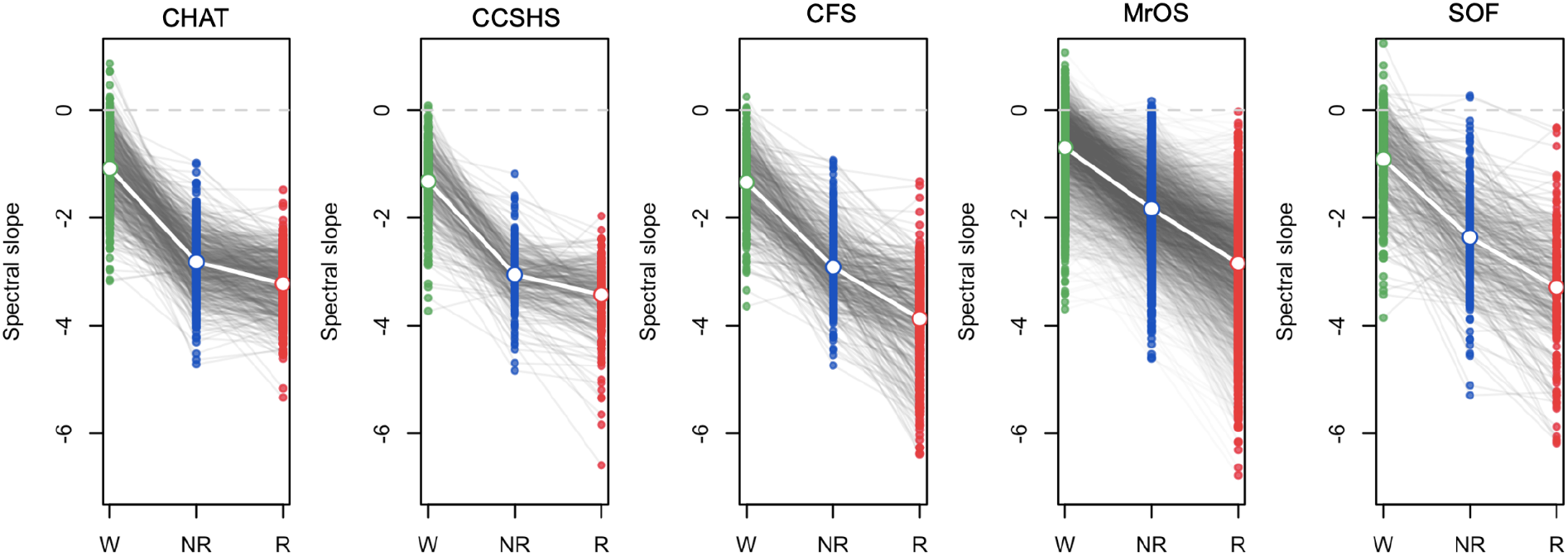
EEG spectral slope for C4-LM. See legend of **Figure 2** for details: this plot provides the same analysis but for C4-LM instead of C3-LM. Green, blue and red indicate wake, NREM and REM respectively. Also see **Table S8**.

**Figure S9.**
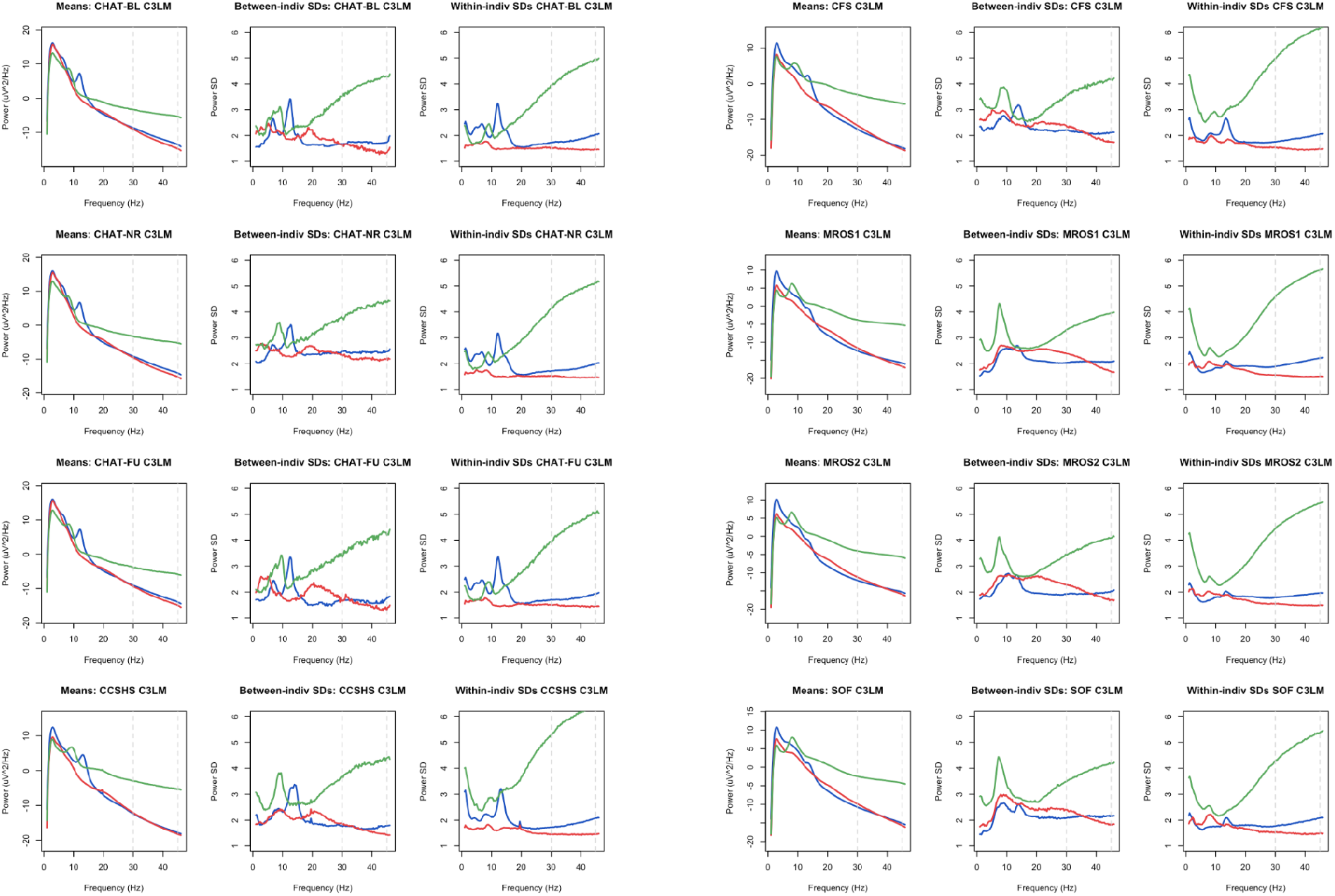
Spectral power means and variability, by sleep state and cohort. See legend of Figure 4 for details: here, figures are plotted separately by cohort rather than super-imposed (as in **Figure 4**). Green, blue and red indicate wake, NREM and REM respectively.

**Figure S10.**
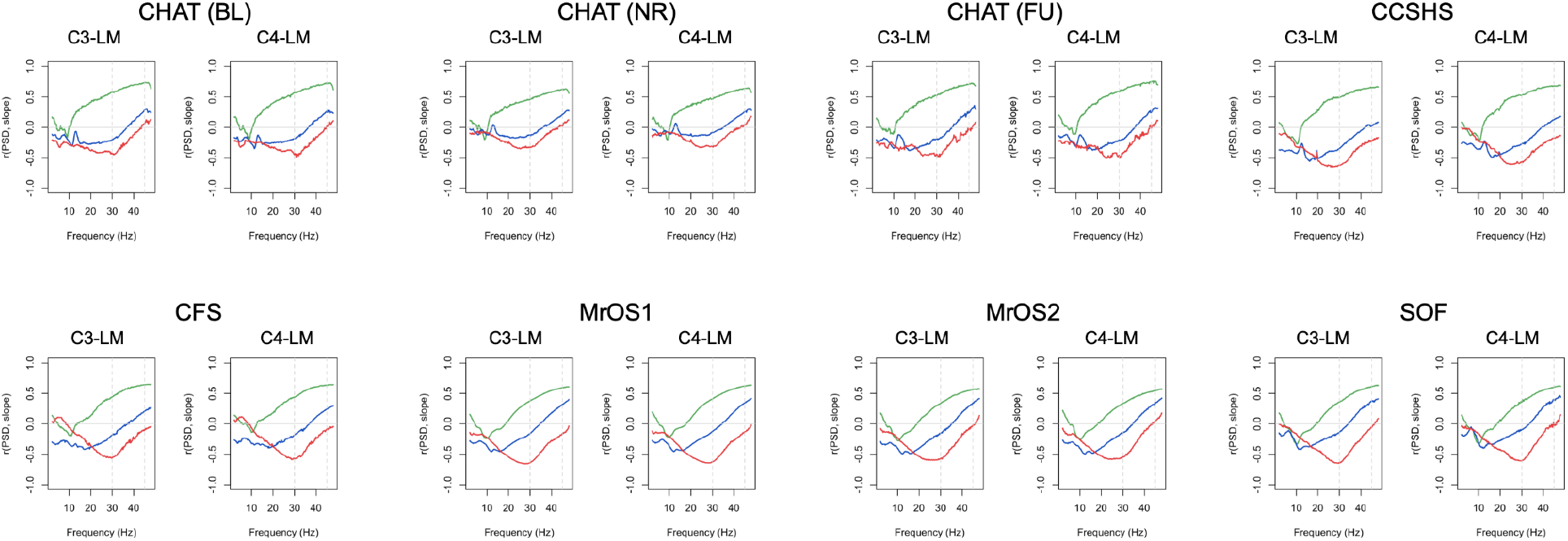
Correlations between EEG spectral slope and power. All analyses based on the LM-referenced dataset. These Figures provide similar information as Figure 6 (top row) in the main text: here, all non-SHHS cohorts are plotted, results for both LM-referenced central channels are given; also, the x-axis extends < 10 Hz whereas **Figure 6** excluded that portion of the power spectrum. Green, blue and red indicate wake, NREM and REM respectively.

**Figure S11.**
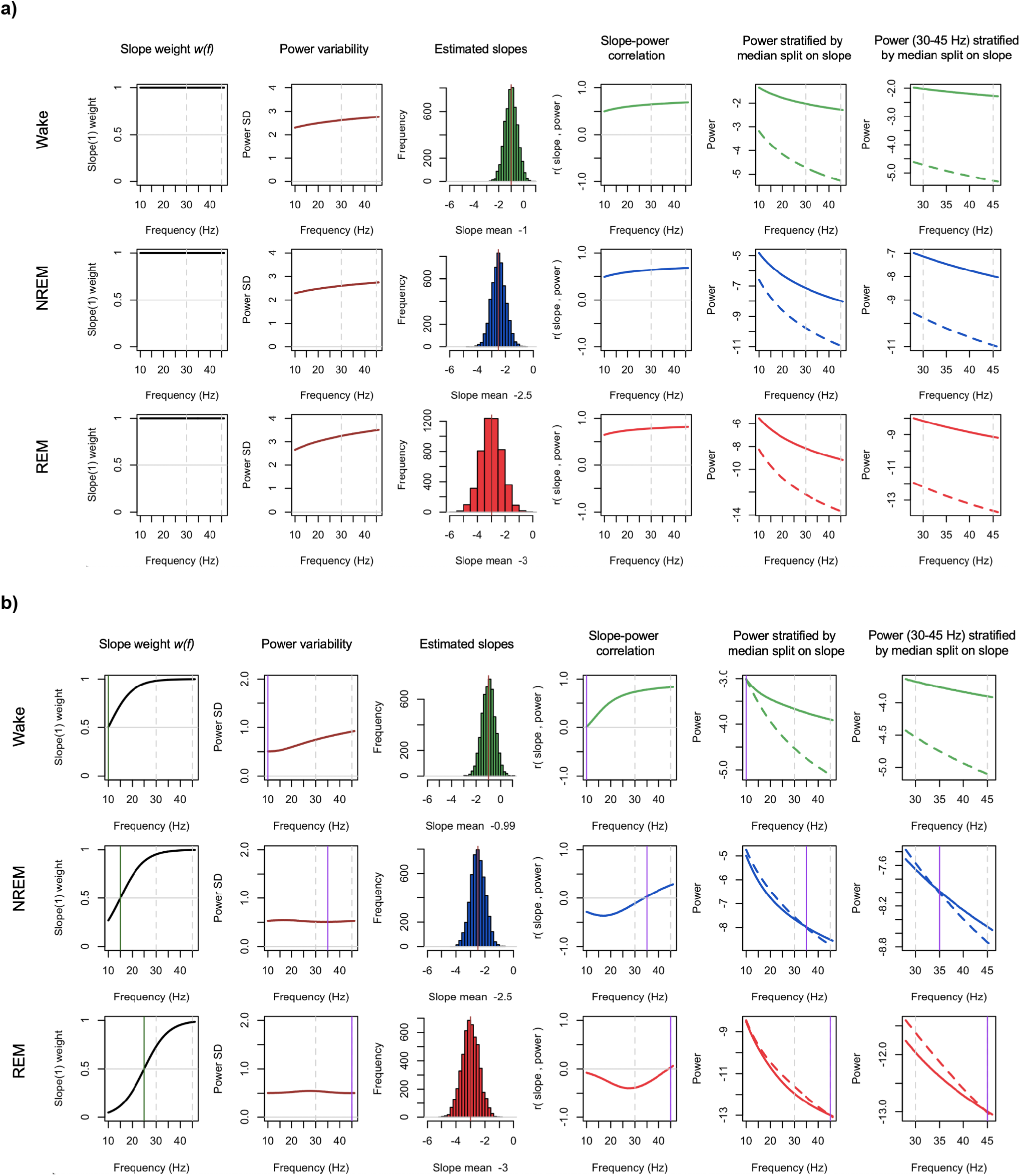
Simulated power spectra with estimated power, spectral slopes, and their correlations. Based on *N* = 5,000 simulated spectra, derived statistics (columns 2 to 6) for a) the original model parameterization, assuming a strict power law model with mean *α* = 1, 2.5 and 3 for wake, NREM and REM respectively (and SDs of 0.5, 0.5 and 0.75, approximately following the observed between-individual estimates from **Figure 5**), and **b)** a revised model, with similar population parameters for slope means and variances but allowing different centers of rotation (*f_r_* = 10, 35 and 45 Hz for wake, NREM and REM respectively) and setting *w*(*f*) such that variation in *α* had less influence on the slope at lower frequencies. Green, blue and red indicate wake, NREM and REM respectively. See **Methods** for details.

**Figure S12.**
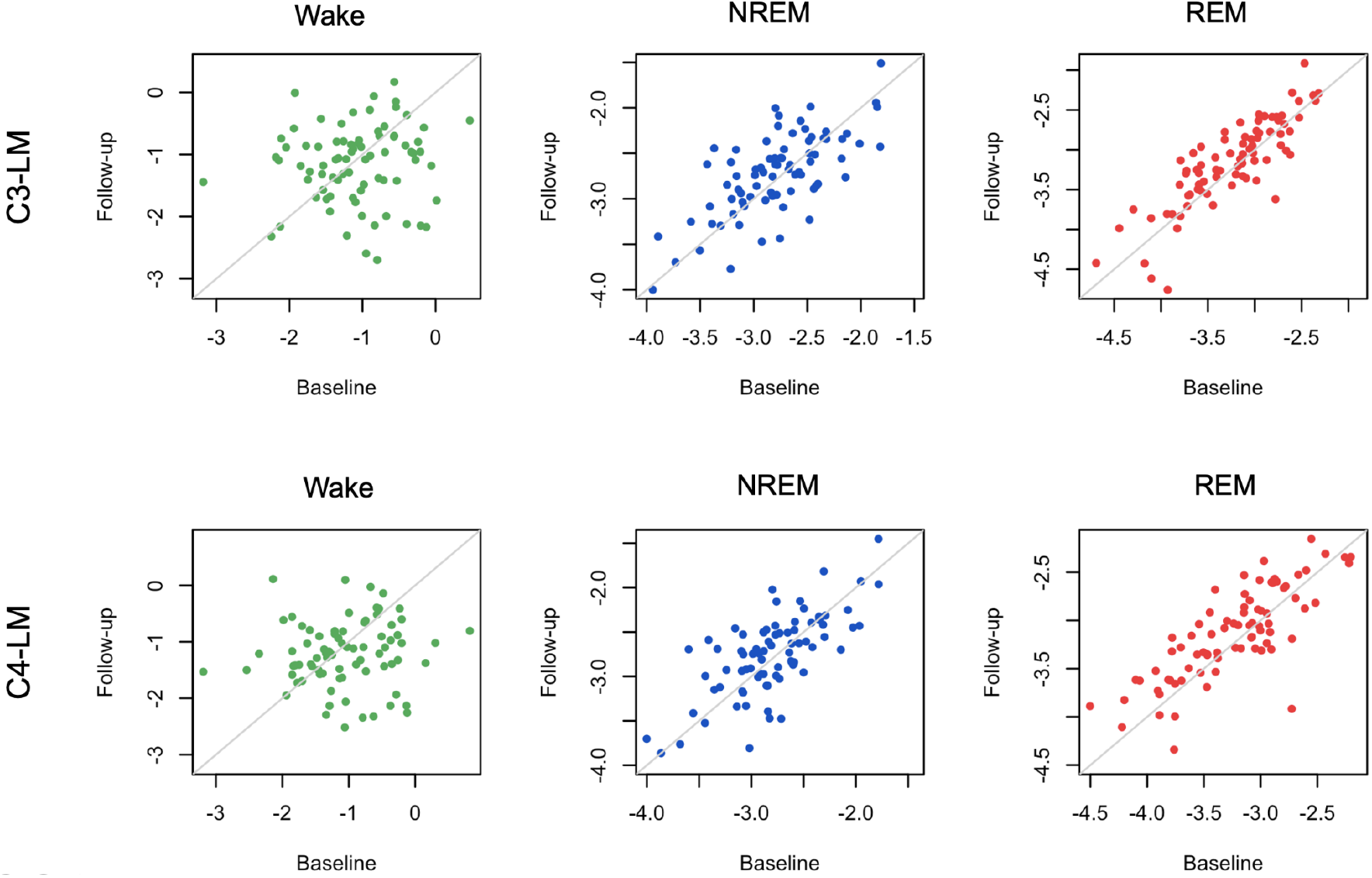
Longitudinal analyses of the CHAT cohort. Based on the LM-referenced dataset, scatter-plots show the EEG spectral slope for baseline and follow-up CHAT (childhood) studies (*N* = 80 pairs post QC, ~6 months interval) stratified by sleep state and channel (C3-LM and C4-LM). Green, blue and red indicate wake, NREM and REM respectively. See **Table S11** for statistical results.

**Table S1.**
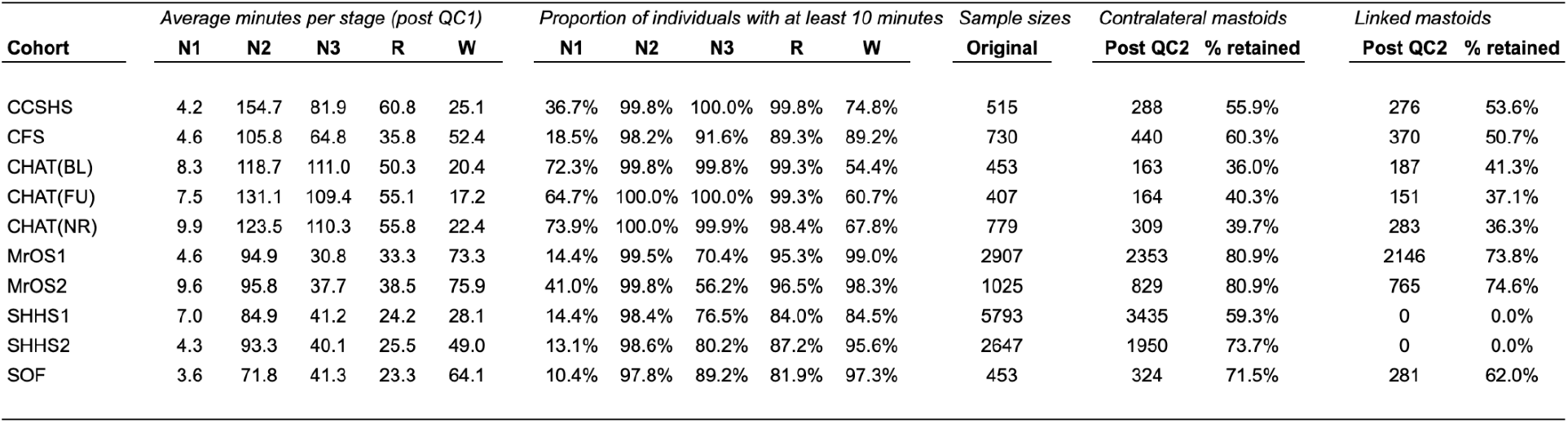
Pre-QC and post-QC sample sizes and durations of sleep state by cohort. For each cohort, the average duration of each state, following the initial round of QC (based on removing individuals without sufficient duration (at least 10 minutes) after performing various epoch-level exclusions (QC1), e.g. only retaining epochs flanked by similarly staged epochs, rejecting epochs with annotated arousals or respiratory events, as well as signal outliers, etc. The second round of QC (QC2) excluded individuals based upon statistical properties of the derived metrics (e.g. power, slope). See **Methods** for details. These procedures were collectively designed to be stringent: that they removed large proportions of some cohorts for the final analysis more reflects the choices of QC rather than inherent issues with the data (i.e. many childhood recordings were removed due to low rates of WASO, as we excluded leading and trailing wake epochs from all recordings, but required all studies to have sufficient duration of wake as well as sleep epochs). No SHHS individuals were retained in the linked mastoid dataset, as it was not possible to re-reference the hardwired contralateral mastoid channels; also see **Supplementary Information, Section 1** for other technical issues encountered in the SHHS cohorts.

**Table S2.** Mean EEG slopes stratified by state and differences between slopes: CM-referenced dataset. See the **Methods** for details on the calculation of spectral slopes.

| Cohort | Channel | Mean slope |  |  | Mean difference |  | t-statistic |  |  |
| --- | --- | --- | --- | --- | --- | --- | --- | --- | --- |
|  |  | W | NR | R | NR - W | R - W | NR - W | R - NR | R - W |
| CHAT | C3-M2 | -1.10 | -2.91 | -3.34 | -1.81 | -2.24 | -53.41 | -19.08 | -65.94 |
|  | C4-M1 | -1.15 | -2.98 | -3.43 | -1.83 | -2.27 | -51.99 | -18.62 | -64.39 |
|  | C3-C4 | -1.73 | -2.81 | -3.08 | -1.08 | -1.36 | -29.72 | -13.37 | -36.15 |
|  | M1-M2 | -0.96 | -3.08 | -3.75 | -2.12 | -2.80 | -49.06 | -21.37 | -64.09 |
| CCSHS | C3-M2 | -1.42 | -3.01 | -3.44 | -1.59 | -2.03 | -42.49 | -14.71 | -48.00 |
|  | C4-M1 | -1.41 | -3.17 | -3.61 | -1.76 | -2.19 | -40.04 | -13.53 | -46.16 |
|  | C3-C4 | -1.50 | -3.07 | -3.23 | -1.56 | -1.73 | -33.52 | -6.09 | -33.21 |
|  | M1-M2 | -1.34 | -2.70 | -3.29 | -1.36 | -1.95 | -24.12 | -15.58 | -32.08 |
| CFS | C3-M2 | -1.51 | -3.12 | -4.03 | -1.62 | -2.52 | -42.78 | -23.76 | -55.04 |
|  | C4-M1 | -1.58 | -3.31 | -4.20 | -1.73 | -2.62 | -39.82 | -22.97 | -56.67 |
|  | C3-C4 | -1.67 | -3.10 | -3.67 | -1.43 | -2.00 | -37.11 | -19.49 | -42.73 |
|  | M1-M2 | -1.45 | -3.12 | -3.85 | -1.67 | -2.40 | -28.39 | -17.65 | -37.07 |
| MrOS | C3-M2 | -0.77 | -1.87 | -2.87 | -1.10 | -2.11 | -75.63 | -68.93 | -120.59 |
|  | C4-M1 | -0.79 | -1.96 | -2.91 | -1.17 | -2.12 | -76.45 | -65.11 | -119.58 |
|  | C3-C4 | -1.09 | -2.42 | -2.88 | -1.33 | -1.79 | -78.59 | -42.75 | -93.89 |
|  | M1-M2 | -0.55 | -1.43 | -2.38 | -0.88 | -1.83 | -47.15 | -65.07 | -85.60 |
| SOF | C3-M2 | -0.97 | -2.32 | -3.24 | -1.35 | -2.27 | -30.81 | -22.01 | -46.88 |
|  | C4-M1 | -0.94 | -2.36 | -3.30 | -1.42 | -2.36 | -30.79 | -22.89 | -47.27 |
|  | C3-C4 | -1.52 | -2.77 | -3.09 | -1.26 | -1.57 | -25.50 | -9.74 | -28.24 |
|  | M1-M2 | -0.48 | -1.58 | -2.68 | -1.10 | -2.20 | -20.40 | -23.82 | -34.43 |

**Table S3.**
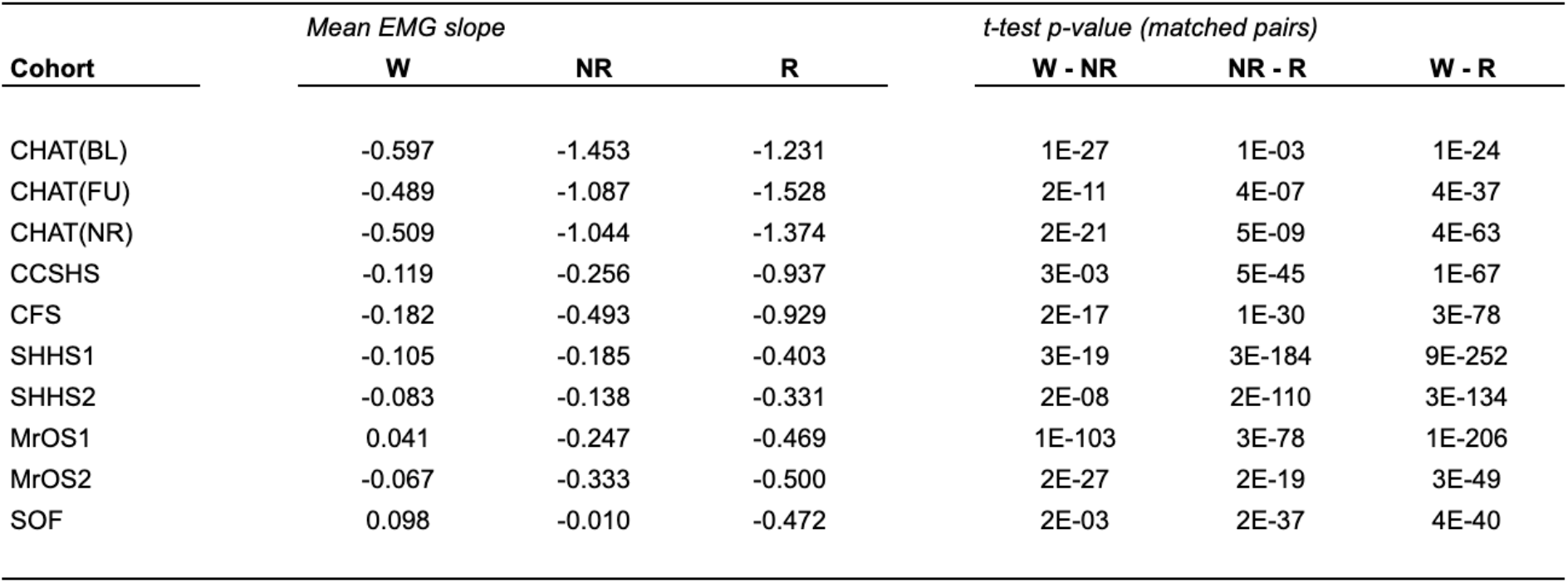
Mean EMG slopes by state and cohort. See **Methods** for details on the calculation of EMG spectral slopes. Also see **Figure S3**.

**Table S4.**
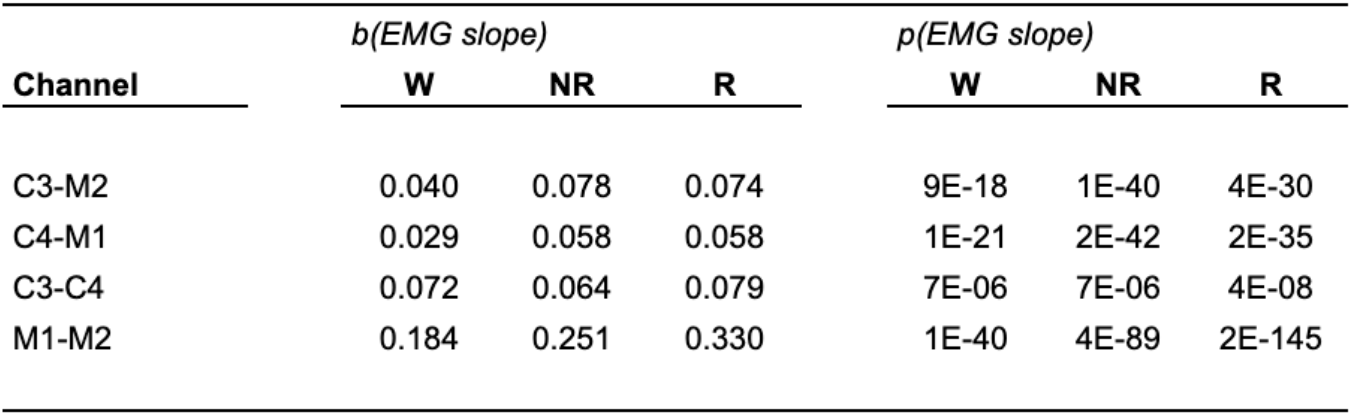
Associations between EEG and EMG spectral slopes in the CM-referenced dataset. Standardized regression coefficients and associated *p*-values from a regression of EEG slope on EMG controlling for age (up to 5th-order polynomials), sex, cohort, race, BMI, AHI and AI. All analyses were performed in the CM-referenced dataset.

**Table S5.**
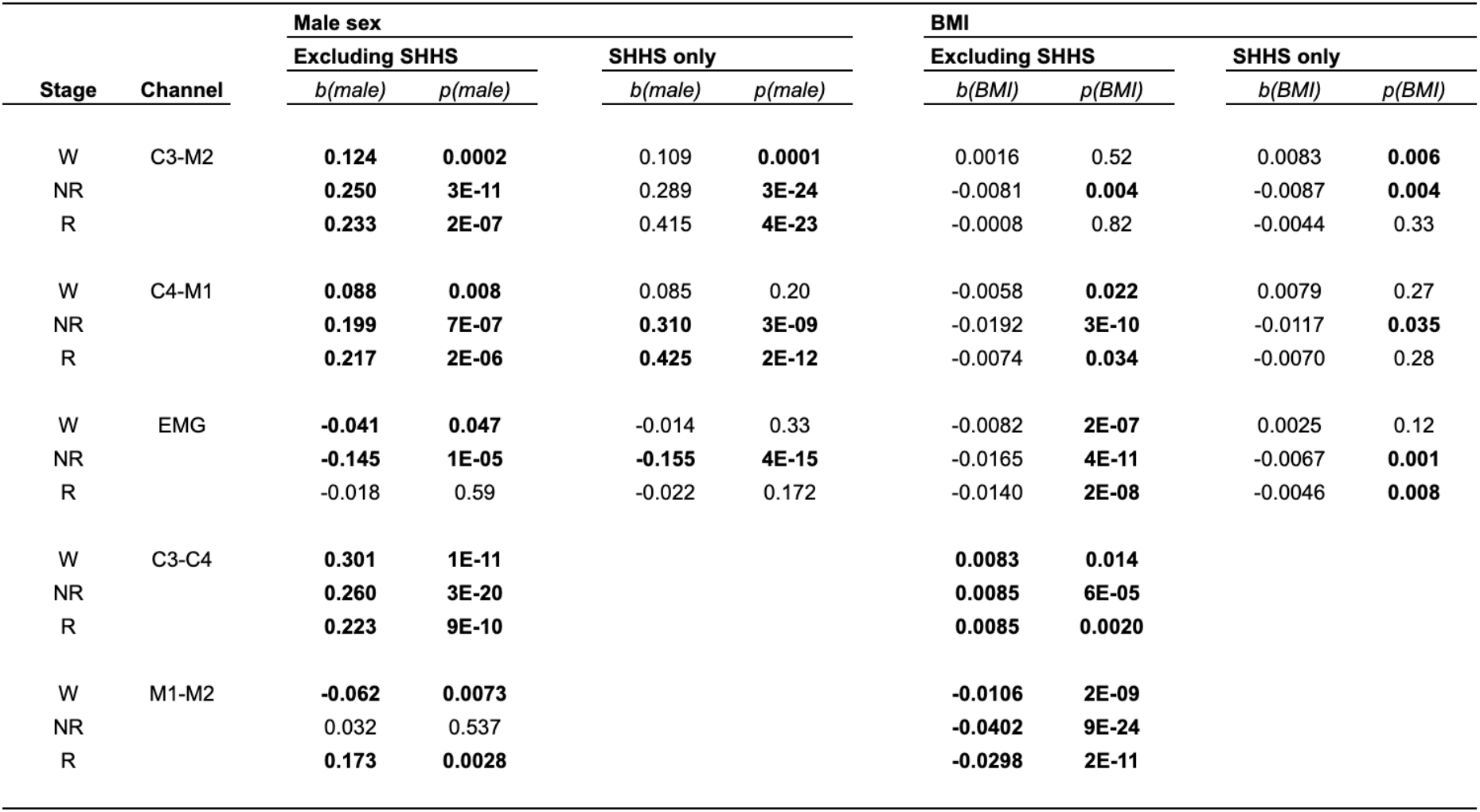
EEG spectral slope associations with sex and BMI in the CM-referenced dataset. Coefficients and *p*-values from linear regression models of slope on sex and BMI, additionally controlling for age (and higher-order terms), race and cohort. Also see **Figure S5**.

| Stage | Channel | Male sex |  |  |  | BMI |  |  |  |
| --- | --- | --- | --- | --- | --- | --- | --- | --- | --- |
|  |  | Excluding SHHS |  | SHHS only |  | Excluding SHHS |  | SHHS only |  |
|  |  | *b*(male) | *p*(male) | *b*(male) | *p*(male) | *b*(BMI) | *p*(BMI) | *b*(BMI) | *p*(BMI) |
| W | C3-M2 | **0.124** | **0.0002** | 0.109 | **0.0001** | 0.0016 | 0.52 | 0.0083 | **0.006** |
| NR |  | **0.250** | **3E-11** | 0.289 | **3E-24** | -0.0081 | **0.004** | -0.0087 | **0.004** |
| R |  | **0.233** | **2E-07** | 0.415 | **4E-23** | -0.0008 | 0.82 | -0.0044 | 0.33 |
| W | C4-M1 | **0.088** | **0.008** | 0.085 | 0.20 | -0.0058 | **0.022** | 0.0079 | 0.27 |
| NR |  | **0.199** | **7E-07** | **0.310** | **3E-09** | -0.0192 | **3E-10** | -0.0117 | **0.035** |
| R |  | **0.217** | **2E-06** | **0.425** | **2E-12** | -0.0074 | **0.034** | -0.0070 | 0.28 |
| W | EMG | **-0.041** | **0.047** | -0.014 | 0.33 | -0.0082 | **2E-07** | 0.0025 | 0.12 |
| NR |  | **-0.145** | **1E-05** | **-0.155** | **4E-15** | -0.0165 | **4E-11** | -0.0067 | **0.001** |
| R |  | -0.018 | 0.59 | -0.022 | 0.172 | -0.0140 | **2E-08** | -0.0046 | **0.008** |
| W | C3-C4 | **0.301** | **1E-11** |  |  | **0.0083** | **0.014** |  |  |
| NR |  | **0.260** | **3E-20** |  |  | **0.0085** | **6E-05** |  |  |
| R |  | **0.223** | **9E-10** |  |  | **0.0085** | **0.0020** |  |  |
| W | M1-M2 | **-0.062** | **0.0073** |  |  | **-0.0106** | **2E-09** |  |  |
| NR |  | 0.032 | 0.537 |  |  | **-0.0402** | **9E-24** |  |  |
| R |  | **0.173** | **0.0028** |  |  | **-0.0298** | **2E-11** |  |  |

**Table S6.** EEG spectral slope associations with sex and BMI in the LM-referenced dataset. Coefficients and *p*-values from linear regression models of slope on sex and BMI, additionally controlling for age (and higher-order terms), race and cohort. The SHHS was excluded from all LM-reference analyses.

| Channel | Stage | Male sex |  | BMI |  |
| --- | --- | --- | --- | --- | --- |
|  |  | *b*(male) | *p*(male) | *b*(BMI) | *p*(BMI) |
| C3-LM | W | **0.178** | **3E-08** | 0.004 | 0.072 |
|  | NR | **0.343** | **5E-25** | 0.005 | 0.07 |
|  | R | **0.318** | **1E-14** | **0.009** | **0.003** |
| C4-LM | W | **0.158** | **1E-06** | 0.002 | 0.46 |
|  | NR | **0.330** | **2E-23** | 0.002 | 0.42 |
|  | R | **0.333** | **1E-15** | **0.008** | **0.009** |
| C3-M2 | W | **0.140** | **4E-06** | 0.002 | 0.41 |
|  | NR | **0.285** | **4E-15** | -0.008 | **0.0061** |
|  | R | **0.286** | **2E-12** | 0.001 | 0.78 |
| C4-M1 | W | **0.109** | **0.0005** | -0.005 | 0.029 |
|  | NR | **0.224** | **9E-09** | -0.017 | **1E-08** |
|  | R | **0.259** | **6E-10** | -0.006 | 0.05 |
| C3-C4 | W | **0.323** | **1E-12** | **0.009** | **0.010** |
|  | NR | **0.276** | **1E-22** | **0.008** | **6E-05** |
|  | R | **0.217** | **9E-10** | **0.008** | **0.002** |
| M1-M2 | W | **-0.056** | **0.02** | **-0.012** | **3E-11** |
|  | NR | 0.016 | 0.77 | **-0.043** | **1E-24** |
|  | R | **0.155** | **0.0106** | **-0.036** | **1E-14** |
| EMG | W | -0.045 | 0.047 | **-0.007** | **1E-05** |
|  | NR | -0.169 | 5E-06 | **-0.015** | **1E-07** |
|  | R | -0.038 | 0.30 | **-0.017** | **1E-09** |

**Table S7.** Associations between EEG and EMG slopes in the LM-referenced dataset. Correlation coefficients and *p*-values for associations between EEG and EMG slopes in the LM-referenced dataset. Equivalent results are also reported for the CM-derived EEG slopes (in this same dataset). EEG-EMG correlations are attenuated comparing LM-derived to CM-derived estimates (albeit still significantly larger than zero). Similar results were obtained when controlling for age, sex, cohort and other covariates (see legend for **Table S4** for details; note: **Table S4** shows standardized regression coefficients from the adjusted model, and so the CM results are not directly comparable to the correlation coefficient presented here).

| Channel | $r(\text{EMG slope})$ | | | $p(\text{EMG slope})$ | | |
| --- | --- | --- | --- | --- | --- | --- |
|  | W | NR | R | W | NR | R |
| C3-LM | 0.114 | 0.092 | 0.045 | 6E-10 | 2E-08 | 2E-02 |
| C4-LM | 0.109 | 0.104 | 0.067 | 2E-09 | 1E-10 | 2E-04 |
| C3-M2 | 0.153 | 0.220 | 0.190 | < 1E-15 | < 1E-15 | < 1E-15 |
| C4-M1 | 0.154 | 0.245 | 0.217 | < 1E-15 | < 1E-15 | < 1E-15 |

**Table S8.** Mean EEG slopes and state differences in the LM-referenced dataset. Comparable state-specific slope means, and tests of state differences as shown in **Table S2** (for the CM-dataset/channels), but here for the LM-derived slopes.

| Cohort | Channel | Mean EEG spectral slope |  |  |  |  | Within-individual, between stage t-statistics |  |  |  |  |
| --- | --- | --- | --- | --- | --- | --- | --- | --- | --- | --- | --- |
|  |  | W | N1 | N2 | N3 | R | NR - W | R - NR | R - W | N2 - N1 | N2 - N3 |
| CHAT (BL+NR) | C3-LM | -1.10 | -2.58 | -2.81 | -2.54 | -3.20 | -54.40 | -18.97 | -65.45 | -8.81 | -13.14 |
|  | C4-LM | -1.12 | -2.58 | -2.82 | -2.56 | -3.21 | -52.59 | -18.64 | -64.98 | -9.19 | -12.48 |
| CCSHS | C3-LM | -1.38 | -2.69 | -3.00 | -2.81 | -3.40 | -44.10 | -13.03 | -47.94 | -6.11 | -8.90 |
|  | C4-LM | -1.38 | -2.69 | -3.05 | -2.87 | -3.44 | -42.32 | -13.14 | -46.44 | -7.13 | -8.58 |
| CFS | C3-LM | -1.38 | -2.72 | -2.84 | -2.65 | -3.77 | -41.99 | -23.76 | -54.28 | -0.71 | -10.27 |
|  | C4-LM | -1.40 | -2.78 | -2.91 | -2.72 | -3.85 | -42.21 | -23.58 | -55.32 | -0.86 | -9.31 |
| MrOS | C3-LM | -0.73 | -1.77 | -1.83 | -1.61 | -2.81 | -77.92 | -69.05 | -119.00 | 2.37 | -24.18 |
|  | C4-LM | -0.73 | -1.77 | -1.84 | -1.63 | -2.80 | -78.99 | -68.43 | -119.30 | 1.62 | -22.80 |
| SOF | C3-LM | -0.96 | -2.22 | -2.35 | -1.99 | -3.24 | -32.52 | -21.10 | -44.61 | 0.21 | -14.03 |
|  | C4-LM | -0.93 | -2.16 | -2.36 | -1.99 | -3.24 | -31.23 | -22.07 | -43.48 | -0.35 | -13.57 |

**Table S9.** Cross-state correlations in EEG spectral slopes. All statistics based on the LM-reference datasets (slopes for C3-LM and C4-LM). Correlations stratified by cohort, with outliers (+/- 3 SD units) removed. All correlations are significantly greater than 0 (*p* < 10^−10^).

| Cohort | Channel | EEG slope cross-stage correlations |  |  |
| --- | --- | --- | --- | --- |
| | | $r(NR, R)$ | $r(R, W)$ | $r(NR, W)$ |
| CCSHS | C3-LM | 0.53 | 0.24 | 0.32 |
|  | C4-LM | 0.55 | 0.22 | 0.31 |
| CFS | C3-LM | 0.51 | 0.43 | 0.46 |
|  | C4-LM | 0.51 | 0.41 | 0.44 |
| CHAT(BL) | C3-LM | 0.62 | 0.28 | 0.28 |
|  | C4-LM | 0.58 | 0.26 | 0.25 |
| CHAT(NR) | C3-LM | 0.67 | 0.21 | 0.24 |
|  | C4-LM | 0.64 | 0.19 | 0.21 |
| CHAT(FU) | C3-LM | 0.58 | 0.41 | 0.43 |
|  | C4-LM | 0.56 | 0.35 | 0.41 |
| MrOS1 | C3-LM | 0.65 | 0.47 | 0.52 |
|  | C4-LM | 0.64 | 0.47 | 0.50 |
| MrOS2 | C3-LM | 0.73 | 0.55 | 0.58 |
|  | C4-LM | 0.76 | 0.55 | 0.57 |
| SOF | C3-LM | 0.61 | 0.44 | 0.52 |
|  | C4-LM | 0.68 | 0.45 | 0.51 |

**Table S10.** Test-retest correlations and mean differences in the EEG spectral slope (MrOS and CHAT). Also see **Figure 10** (MrOS) and **Figure S12** (CHAT) for plots of test-retest EEG slope distributions.

| Cohort | Channel | Stage(s) | Test-retest correlations |  | Mean EEG slopes |  | Test-retest mean differences |  |
| --- | --- | --- | --- | --- | --- | --- | --- | --- |
|  |  |  | r | p-value | Baseline | Follow-up | Difference | p-value |
| **MrOS**<br>(n=610 pairs, post QC)<br>~6 years | C3-LM | W | 0.55 | 8E-48 | -0.7 | -0.75 | -0.04 | 0.08 |
|  |  | NR | 0.69 | 3E-87 | -1.84 | -1.67 | 0.17 | 1E-15 |
|  |  | R | 0.80 | 8E-138 | -2.89 | -2.53 | 0.37 | 4E-50 |
|  | C4-LM | W | 0.51 | 7E-41 | -0.709 | -0.709 | 0.00 | 0.98 |
|  |  | NR | 0.63 | 5E-68 | -1.84 | -1.53 | 0.32 | 4E-42 |
|  |  | R | 0.75 | 6E-108 | -2.86 | -2.3 | 0.57 | 5E-87 |
|  | C3-LM | R - NR | 0.67 | 2E-78 | -1.04 | -0.853 | 0.20 | 9E-20 |
|  |  | R - W | 0.62 | 9E-64 | -2.17 | -1.76 | 0.41 | 1E-39 |
|  |  | NR - W | 0.49 | 2E-37 | -1.13 | -0.908 | 0.22 | 3E-16 |
|  | C4-LM | R - NR | 0.67 | 3E-80 | -1.01 | -0.778 | 0.25 | 2E-31 |
|  |  | R - W | 0.55 | 1E-48 | -2.14 | -1.58 | 0.57 | 3E-63 |
|  |  | NR - W | 0.43 | 1E-28 | -1.13 | -0.801 | 0.32 | 4E-30 |
| Cohort | Channel | Stage(s) | r | p-value | Baseline | Follow-up | Difference | p-value |
| **CHAT**<br>(n=80 pairs, post QC)<br>~6 months | C3-LM | W | 0.10 | 0.38 | -1.08 | -1.15 | -0.08 | 0.42 |
|  |  | N2 | 0.69 | 1E-12 | -2.79 | -2.72 | 0.07 | 0.1 |
|  |  | R | 0.84 | 9E-23 | -3.26 | -3.16 | 0.09 | 0.006 |
|  | C4-LM | W | 0.08 | 0.46 | -1.06 | -1.21 | -0.16 | 0.1 |
|  |  | N2 | 0.71 | 4E-13 | -2.81 | -2.75 | 0.05 | 0.2 |
|  |  | R | 0.79 | 4E-18 | -3.26 | -3.14 | 0.10 | 0.004 |
|  | C3-LM | R - NR | 0.71 | 3E-13 | -0.456 | -0.431 | 0.03 | 0.52 |
|  |  | R - W | 0.28 | 0.01 | -2.17 | -2.02 | 0.17 | 0.07 |
|  |  | NR - W | 0.20 | 0.08 | -1.72 | -1.57 | 0.16 | 0.08 |
|  | C4-LM | R - NR | 0.58 | 2E-08 | -0.441 | -0.371 | 0.05 | 0.21 |
|  |  | R - W | 0.26 | 0.021 | -2.19 | -1.92 | 0.27 | 0.005 |
|  |  | NR - W | 0.23 | 0.041 | -1.76 | -1.55 | 0.22 | 0.02 |

**Table S11.** Cross-sectional analysis of age-related flattening of the spectral slope. All statistics are based on the LM-reference datasets, using a multiple linear regression model of EEG slope (here average of C3-LM and C4-LM) on age (linear) plus covariates, performed within cohort. CCSHS was excluded as there was effectively no variation in age (most participants were either 17 or 18 years of age). Similar patterns of results were obtained for analyses of each individual LM-referenced channel.

| **Cohort** | **Wake** |  |  | **NR** |  |  | **R** |  |  |
| --- | --- | --- | --- | --- | --- | --- | --- | --- | --- |
|  | *Mean* | *b(age)* | *p(age)* | *Mean* | *b(age)* | *p(age)* | *Mean* | *b(age)* | *p(age)* |
| CHAT(BL) | -1.11 | 0.071 | 0.058 | -2.81 | **0.158** | **6E-08** | -3.23 | **0.134** | **3E-06** |
| CHAT(NR) | -1.07 | **0.081** | **0.002** | -2.82 | **0.135** | **3E-08** | -3.17 | **0.117** | **4E-07** |
| CFS | -1.37 | -0.001 | 0.68 | -2.87 | **0.005** | **0.006** | -3.85 | **-0.009** | **1E-04** |
| MrOS1 | -0.71 | 0.001 | 0.69 | -1.82 | -0.002 | 0.53 | -2.85 | **0.009** | **0.0081** |
| SOF | -0.94 | -0.003 | 0.82 | -2.34 | -0.002 | 0.88 | -3.25 | 0.017 | 0.32 |

| **Cohort** | **NR-W** |  |  | **R-NR** |  |  | **R-W** |  |  |
| --- | --- | --- | --- | --- | --- | --- | --- | --- | --- |
|  | *Mean* | *b(age)* | *p(age)* | *Mean* | *b(age)* | *p(age)* | *Mean* | *b(age)* | *p(age)* |
| CHAT(BL) | -1.70 | **0.091** | **0.033** | -0.40 | -0.024 | 0.39 | -2.10 | 0.049 | 0.27 |
| CHAT(NR) | -1.75 | 0.054 | 0.094 | -0.35 | -0.018 | 0.38 | -2.11 | 0.035 | 0.259 |
| CFS | -1.49 | **0.006** | **0.0025** | -0.99 | **-0.015** | **8E-13** | -2.49 | **-0.009** | **0.00028** |
| MrOS1 | -1.11 | -0.003 | 0.18 | -1.02 | **0.011** | **0.00004** | -2.12 | **0.008** | **0.012** |
| SOF | -1.41 | 0.004 | 0.77 | -0.89 | 0.025 | 0.068 | -2.33 | 0.026 | 0.142 |

